# Chemical ecology of an apex predator life cycle

**DOI:** 10.1101/2021.02.13.430781

**Authors:** Nicholas C. Mucci, Katarina A. Jones, Mengyi Cao, Michael R. Wyatt, Shane Foye, Sarah Kauffman, Michela Taufer, Yoshito Chikaraishi, Shawn Steffan, Shawn Campagna, Heidi Goodrich-Blair

## Abstract

Microbial symbiotic interactions, mediated by small molecule signaling, drive physiological processes of higher order systems. Metabolic analytic technologies advancements provide new avenues to examine how chemical ecology, or conversion of existing biomass to new forms, changes over a symbiotic lifecycle. We examine such processes using the tripartite relationship between nematode host *Steinernema carpocapsae*, its obligate mutualist bacterium, *Xenorhabdus nematophila*, and the insects they infect together. We integrate trophic, metabolomics, and gene regulation analyses to understand insect biomass conversion to nematode or bacterium biomass. Trophic analysis established bacteria as the primary insect consumers, with nematodes at trophic position 4.37, indicating consumption of bacteria and likely other nematodes. Significant, discrete metabolic phases were distinguishable from each other, indicating the insect chemical environment changes reproducibly during bioconversion. Tricarboxylic acid cycle components and amino acids were significantly affected throughout infection. These findings contribute to an ongoing understanding of how symbiont associations shape chemical environments.

**Teaser:** Entomopathogenic nematodes act as an apex predator in some ecosystems through altering chemical environments of their prey.

## Introduction

Symbiotic interactions are ubiquitous in biological systems and have shaped the evolution of life (1). These long-term, intimate associations are driven by small molecule signaling between partners. Bacterial populations establish diverse and expansive metabolite-mediated signaling networks that control gene expression and downstream behaviors, such as biofilm formation and the production of host-interacting effectors (2-4). A common mechanism by which bacteria sense and transduce metabolic signals is through transcription factors whose DNA binding affinity or specificity is modulated by binding metabolite ligands. For instance, LysR-type transcription factors, which are conserved across proteobacteria, are characterized by a conserved *N*-terminal DNA-binding domain and a *C*-terminal domain that varies among LysR-type regulator homologs. The latter domain is responsible for ligand metabolite binding and dictates the response specificity of the transcription factor (5, 6). In a range of bacteria, LysR-type regulators modulate various phenotypes, including virulence, nutrient uptake and metabolic homeostasis, motility, quorum sensing, and antibiotic resistance (7). Another diverse family of transcription factors, feast/famine regulatory factors like leucine-responsive regulatory protein (Lrp), will bind and dimerize with amino acids in response to nutrient levels and globally induce transcriptional changes (8).

Given the key function of metabolites in communicating information about intracellular and extracellular environmental conditions, examining their identities and abundances is critical to understanding biological systems. Metabolomics has enabled such studies and has been used to detect specific small molecules that drive essential cellular processes and inter-kingdom signaling (9, 10). Further, it is being applied to more complex ecosystems comprising multi-species microbiota colonizing a host (11). However, to date such studies have been primarily focused on binary conditional comparisons between treatments, or on single, snapshot sampling of complex interactions. Here, to gain insights temporal changes in metabolic pathways that occur in complex ecosystems, a longitudinal analysis of metabolic profiles was conducted in closed ecosystem in which biomass is being reproducibly converted from one type of living organism to another. The closed ecosystem comprised an individual insect infected with an entomopathogenic nematode (EPN) and bacterium (EPNB) pair.

EPNs of the genera *Steinernema* and *Heterorhabditis* associate with mutualistic bacteria in the genera *Xenorhabdus* and *Photorhabdus*, respectively. An infective juvenile (IJ) stage of EPN carry their mutualistic bacteria in their intestine as they dwell in the soil seeking insect hosts to infect. Upon infection, the bacteria are released into the insect blood cavity and together the nematode and bacterium kill and consume the insect for their own reproduction before developing into the bacteria-colonized infective stage again to repeat the cycle (12, 13). EPNBs have been applied as insecticide alternatives to promote agricultural productivity and to help prevent transmission of insect diseases like dengue and West Nile virus (14, 15).

In this study, consumption and bioconversion of the insect *Galleria mellonella* by the EPN *Steinernema carpocapsae* and its mutualistic bacterial symbiont, *Xenorhabdus nematophila* was examined from a metabolic perspective. *G. mellonella* is used for laboratory isolation and propagation of EPNB, is a model host to understand virulence of a variety of microbial pathogens and has a characterized metabolome (16). The *S. carpocapsae-X. nematophila* pair was chosen due to the wealth of information available about them from molecular, cellular, and genetic studies (13). For example, it is known that *X. nematophila* bacterial effectors and natural products suppress insect immunity, kill insect blood cells, degrade insect tissues, and defend the insect cadaver from opportunistic competitors (17). Also, *X. nematophila* bacteria are essential for *S. carpocapsae* reproduction; in the absence of bacteria fewer nematode IJs emerge from insect cadavers after reproduction (18). Expression of effectors and physiological adaptation to changing host environments is controlled in *X. nematophila* by transcriptional regulators that are predicted to sense and respond to prevailing metabolic conditions (19). For instance, the LysR-type regulator LrhA that is necessary for *X. nematophila* virulence and controls expression of an extracellular phospholipase that is necessary for insect degradation (12, 19, 20), the sigma factor RpoS that is necessary for colonizing the IJ stage of the nematode (21), the two-component system CpxRA, and the leucine-responsive regulatory protein Lrp, both of which are necessary for normal virulence and mutualism behaviors (17, 22, 23), NilR, a lambda like repressor family transcription factor that negatively regulates genes necessary for nematode colonization (24) and the two-component system OmpR/EnvZ that negatively controls *X. nematophila* swarming motility behavior and exoenzyme production (25). Further, *X. nematophila* displays phenotypic heterogeneity with respect to behaviors important for adaptation to host environments. For instance, “primary form” [1°] *X. nematophila* can be distinguished from “secondary form” [2°] by its motility, antibiotic and natural products secretion, and hemolytic and lipolytic activities and additional phenotypic variants arise over the course of insect infection (26, 27).

The goal of this study was to begin to understand the overall metabolic transformations, or bioconversion processes occurring within a closed yet complex biological ecosystem. ^15^N isotopic enrichment analyses were performed to establish the relative trophic positions of the insect, *G. mellonella*, the nematode, *S. carpocapsae*, and the bacterium, *X. nematophila*, so that the relative roles in bioconversion of each ecosystem member could be established. Then, a metabolomics analysis using an ultra-high performance liquid chromatography high-resolution mass spectrometry (UHPLC-HRMS) metabolomics technique was conducted over a 16-day time course after *S. carpocapsae-X. nematophila* infection of *G. mellonella*, encompassing a complete bioconversion of insect tissues to the bacterial-colonized progeny IJs that emerged from the insect.

## Results

### Trophic analysis reveals *S. carpocapsae* nematodes directly feed on *X. nematophila* bacteria

The trophic identities of the entomopathogenic nematode (*Steinernema carpocapsae*), its bacterial symbiont (*Xenorhabdus nematophila*), and their host insect were measured empirically based on ^15^N isotopic enrichment of amino acids (Table 1 and Data S1). To this end, it was first necessary to establish that the degree of ^15^N-enrichment between the consumers (nematodes, bacteria) and their respective diets (e.g., agar growth media, bacteria, or the insect) was consistent with past studies of inter-trophic enrichment (Supplementary Text). These findings allowed for the subsequent *in vivo* trials involving insect cadavers (Fig. 1B, Table 1 and Data S1). As reported above, the mean TP_glu-phe_ of an uncolonized insect cadaver was 2.2 ± 0.02 (*N* = 6). When the insect was colonized by bacteria alone, the TP_glu-phe_ of the insect-bacteria complex was 2.5 ± 0.03 (*N* = 3). This complex represented the blending of consumer and diet (as described in 28), wherein the consumer (i.e., the *Xenorhabdus* bacterial population) was suffused within and throughout its diet (the insect cadaver). Given that both the bacterial and insect biomass were available within the cadaver, this established the basis for the question as to what a developing nematode would consume/assimilate within the cadaver. The diet of the nematodes (i.e., the insect-bacterial complex) was measured at ∼2.5, thus if the nematodes within the cadaver fed randomly on all available substrates, the nematode TP_glu-phe_ would be expected to be ∼3.5 (i.e., ∼2.5 + 1.0). However, the infective juveniles emerging from the cadavers registered a TP_glu-phe_ of 4.6 ± 0.08 (*N* = 5), a full trophic level higher than expected.

**Table 1.**
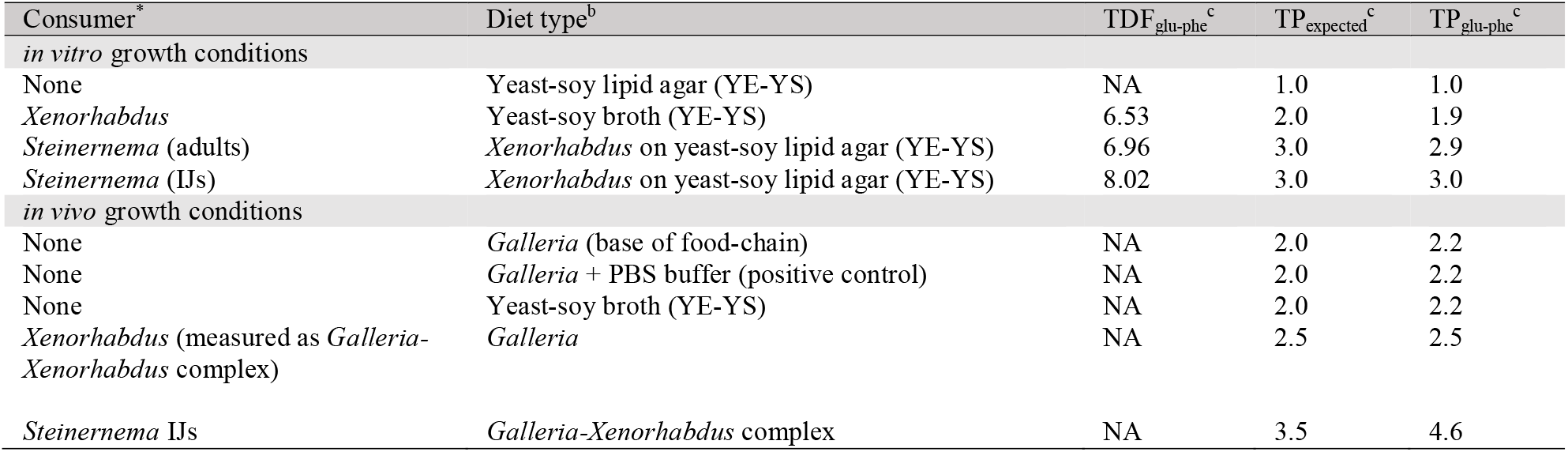
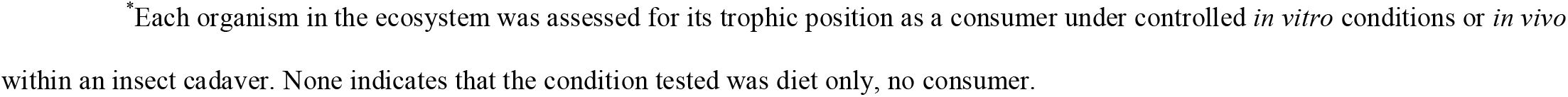
Summary of *in vitro* and *in vivo* trophic measurements.

| Consumer<sup>a</sup> | Diet type<sup>b</sup> | TDF<sub>glu-phe</sub><sup>c</sup> | TP<sub>expected</sub><sup>c</sup> | TP<sub>glu-phe</sub><sup>c</sup> |
| --- | --- | --- | --- | --- |
| *in vitro* growth conditions |  |  |  |  |
| None | Yeast-soy lipid agar (YE-YS) | NA | 1.0 | 1.0 |
| *Xenorhabdus* | Yeast-soy broth (YE-YS) | 6.53 | 2.0 | 1.9 |
| *Steinernema* (adults) | *Xenorhabdus* on yeast-soy lipid agar (YE-YS) | 6.96 | 3.0 | 2.9 |
| *Steinernema* (IJs) | *Xenorhabdus* on yeast-soy lipid agar (YE-YS) | 8.02 | 3.0 | 3.0 |
| *in vivo* growth conditions |  |  |  |  |
| None | *Galleria* (base of food-chain) | NA | 2.0 | 2.2 |
| None | *Galleria* + PBS buffer (positive control) | NA | 2.0 | 2.2 |
| None | Yeast-soy broth (YE-YS) | NA | 2.0 | 2.2 |
| *Xenorhabdus* (measured as *Galleria*-*Xenorhabdus* complex) | *Galleria* | NA | 2.5 | 2.5 |
| *Steinernema* IJs | *Galleria*-*Xenorhabdus* complex | NA | 3.5 | 4.6 |

\*Each organism in the ecosystem was assessed for its trophic position as a consumer under controlled *in vitro* conditions or *in vivo* within an insect cadaver. None indicates that the condition tested was diet only, no consumer.

**Figure 1:**
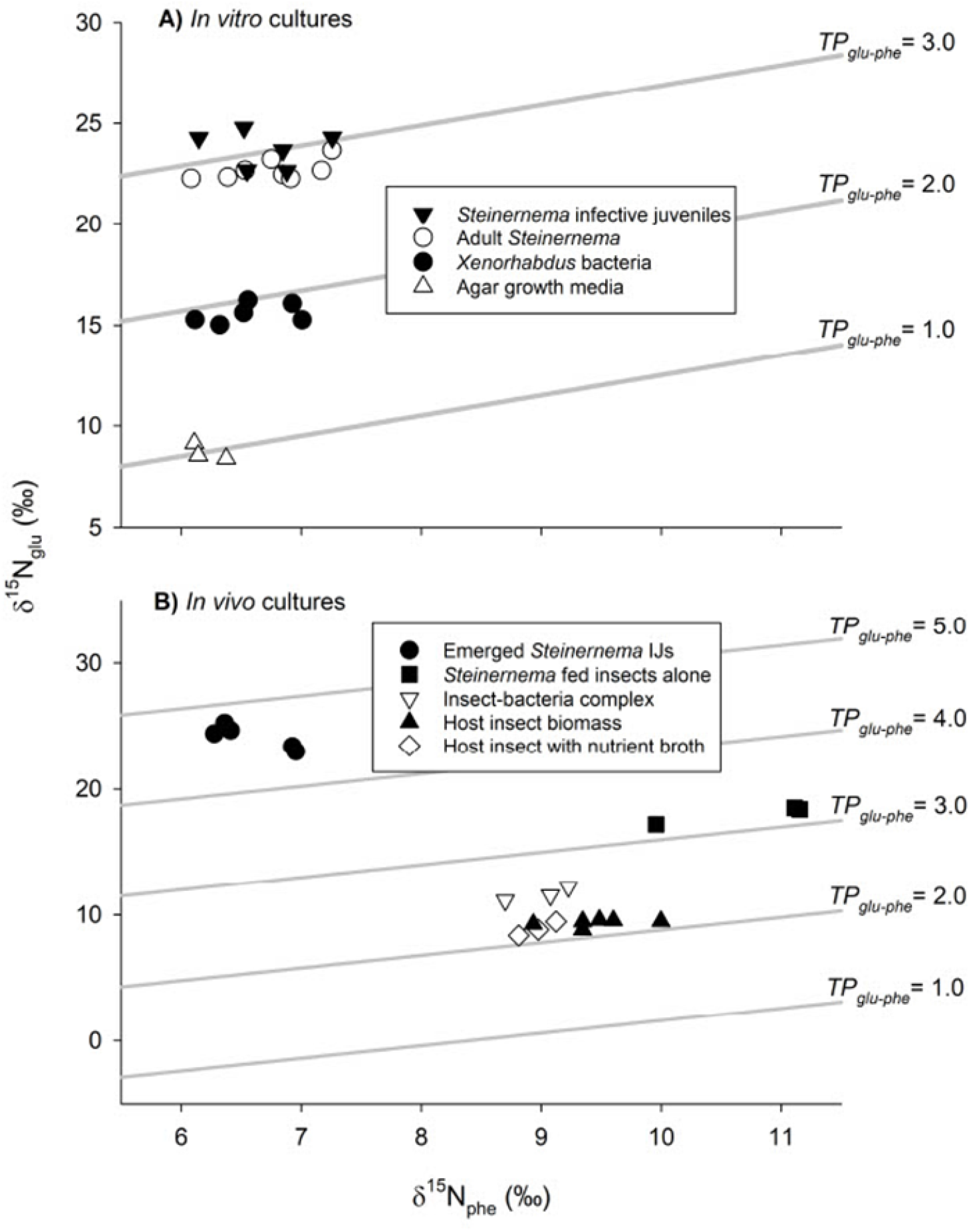
Trophic analyses reveal *Steinernema* nematodes feed on *Xenorhabdus* bacteria (A) *In vitro* and (B) *In vivo*. Trophic isoclines are represented via numeric TP_glu-phe_ % ratios. Specific bacterial cultures or animals are displayed as the different shapes shown in the figure legends.

The data above demonstrated that adult *Steinernema* nematodes consume their mutualistic bacteria. During this stage, *Xenorhabdus* bacteria colonize the anterior intestinal caecum. To determine if this colonization influences the ability of *Steinernema* nematodes to consume *Xenorhabdus*, the TP_glu-phe_ of adult *Steinernema* cultivated on lawns of either wild-type *Xenorhabdus*, or a non-colonizing mutant (Δ*SR1*) was assessed (29). Nematodes had the same TP_glu-phe_ regardless of the colonization proficiency of the bacterial diet, indicating that colonization is not required for nematode direct feeding on its symbiotic bacteria (Data S1).

### *X. nematophila* transcriptional control of metabolic pathways

The trophic analyses described above establish *X. nematophila* bacteria as the linchpin organism in the closed ecosystem, responsible for direct consumption of the insect tissue and serving as a primary food source for its mutualistic host *S. carpocapsae*. To gain insights into the metabolic pathways utilized by *X. nematophila* in performance of these functions, the global regulons of several transcription factors was determined using an exploratory microarray analysis, portions of which have been reported elsewhere (Fig. 2 and Table S1, and Data S2) (30-32). Microarray analyses were conducted on mutants lacking genes encoding the transcription factors LrhA, RpoS, NilR, and Lrp, each of which has a defect in one or more aspects of the *X. nematophila* life cycle (19, 21, 23, 24). In addition, since the primary to secondary form phenotypic variation globally influences host-interaction phenotypes, the transcriptional profiles of these variants were examined from a metabolic perspective. The mutant and secondary form cells were each compared to their wild-type parent or primary form, respectively, using a 2<|fold change| significance cutoff for differences in transcript levels.

**Figure 2:**
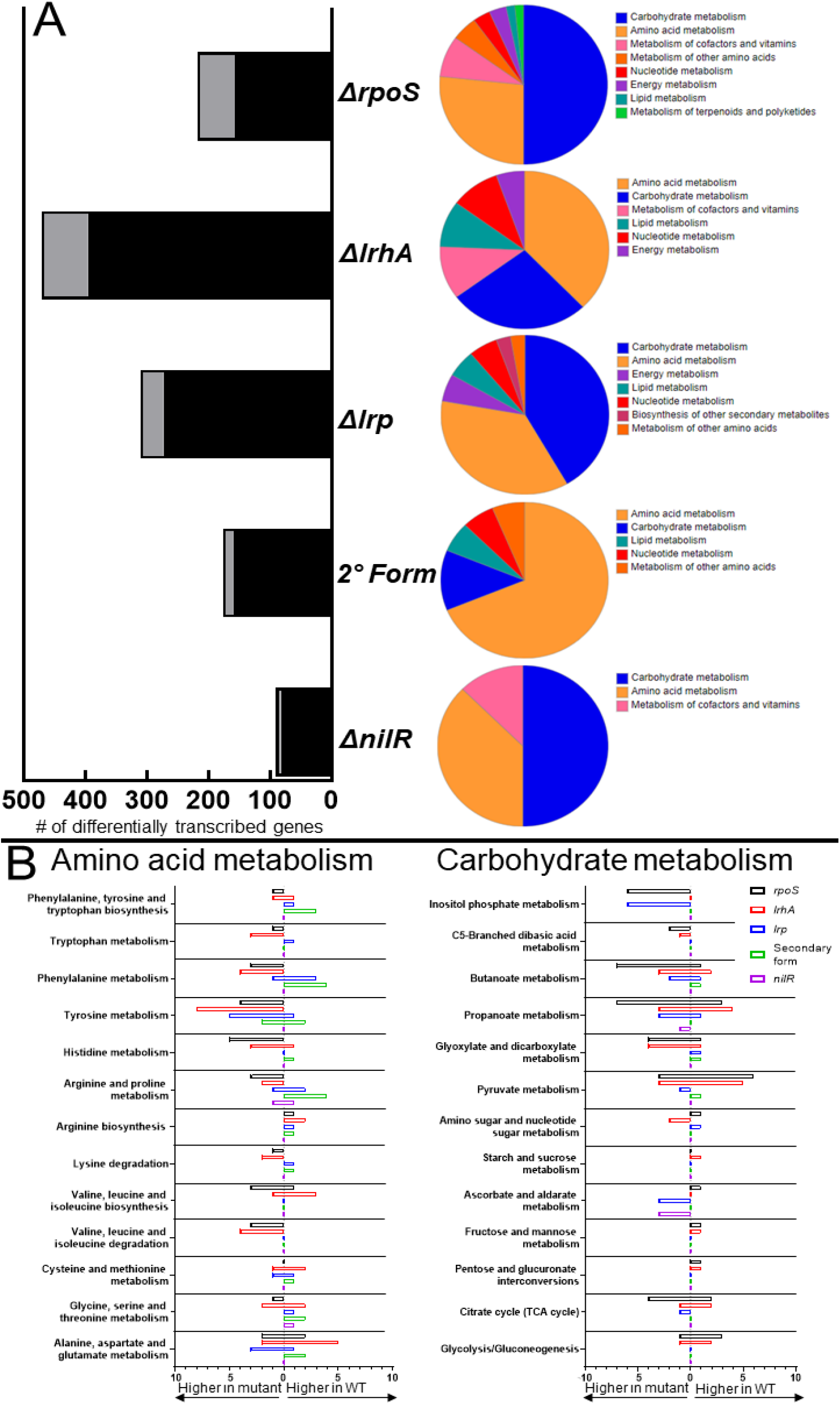
Differentially expressed transcripts between *X. nematophila* mutants and their broad functional categorization. A) Quantification of the number of differentially regulated transcripts and how many are considered metabolic (light grey), as determined by KEGG annotation. |Signal fold change| > 2 was used as a cutoff for significance. BlastKOALA functional categorization of the differential metabolic transcripts are adjacent. The color legend is organized by having the most common category listed first. B) Breakdown of the specific amino acid and carbohydrate metabolism pathways that were affected by the mutations, compared among each strain, with # of genes listed. Positive genes represent transcripts higher in the mutant relative to WT, negative genes represent transcripts lower in the mutant relative to WT.

The number of differential transcripts are a fraction of the 3733 averaged total expressed chromosome ORFs among the strains. The number of genes with differential transcript abundance in mutant/secondary form compared to wild-type/primary was highest in Δ*lrhA* at 396 genes (10.6%), followed by Δ*lrp*, secondary form, Δ*rpoS*, and Δ*nilR* with 273 (7.3%), 159 (4.3%), 157 (4.2%), and 83 (2.2%) differentially abundant transcripts, respectively (Fig. 2A). The proportion of genes categorized as being involved in metabolic activity varied amongst the strains. Through KEGG annotation, the highest proportion of differentially expressed genes categorized as metabolic was observed in the Δ*rpoS* strain, at 38.9%, with Δ*lrhA*, Δ*lrp*, secondary form, and Δ*nilR* having metabolic-related activities at 18.9%, 13.6%, 10.7%, and 9.8%, of the differentially expressed genes in the respective strain. Differential transcript overlap was observed while comparing the 5 strains (Fig. S1). The largest overlap between 2 strains was for the secondary form and Δ*lrp* mutant, consisting of mostly amino acid (*xncB*, the aminotransferase XNC1_2154) and lipid biosynthesis (*fabG, xncL*) genes. *X. nematophila* lacking *lrp* are phenotypically secondary form (23). Another large overlap was observed between Δ*lrp* and Δ*nilR*, which synergistically repress nematode colonization (24). The overlap regulation includes the phosphotransferase system/ascorbate metabolic (XNC1_2826-2828) genes and prokaryotic defense system genes (XNC1_3717-3719, 3724, and 3931).

Functional analysis of the differentially expressed metabolic transcripts was performed through KEGG annotation using BlastKOALA (KEGG Orthology and Links Annotation). Sequences were aligned against a nonredundant set of prokaryotic KEGG genes using BLAST searches (33). Consistent with the KEGG annotation analysis noted above, of the strains tested Δ*rpoS*, Δ*lrhA*, Δ*lrp* were the most strongly impacted with respect to metabolic pathway transcripts. These three strains displayed differences in carbohydrate and amino acid metabolic regulation (Fig. 2A). Branches of carbohydrate metabolism, like propanoate, pentose and glucuronate, and glyoxylate metabolism were impacted by the Δ*lrhA* and Δ*rpoS* mutations (Fig. 2B). Inositol phosphate metabolism was impacted in Δ*rpoS* and Δ*lrp* strains, while pyruvate metabolism was impacted in the Δ*lrhA* strain. Butanoate metabolism, a branch of carbohydrate metabolism where the amino acid ornithine is converted into short-chain fatty acids, was commonly interrupted for all mutant strains except Δ*nilR*. Amino acid biosynthetic pathway transcripts were differently regulated between these three strains, with tyrosine, and the alanine, aspartate, and glutamate biosynthetic pathways similarly disrupted. Histidine and valine metabolism was uniquely altered by the Δ*rpoS* mutation, glycine, serine, and threonine metabolism was uniquely altered by the Δ*lrhA* mutation, and phenylalanine metabolism was uniquely altered by the Δ*lrp* mutation. To investigate the impacts these pathways and others have on the *Xenorhabdus*-*Steinernema* lifecycle, a time course metabolomics experiment was designed to measure the relative quantities of metabolites within them.

### The metabolomic profile of EPNB-infected *G. mellonella*: an overview

Having established that *X. nematophila* bacteria consume infected insect tissue, and in turn the bacteria are consumed by reproducing and developing nematodes, the temporal dynamics of metabolic profiles associated with these processes were examined. *G. mellonella* were infected with *S. carpocapsae* infective juveniles colonized by *X. nematophila* bacteria. Weights of the whole insect samples were relatively similar (Table S2). Insects were sampled over a 16-day time-course. As expected, insects began to die by Hour 24 after infection, and both living and dead insects were sampled at that time point. *X. nematophila* will have reproduced rapidly in order to suppress the insect immune response and release toxins to kill the insect host. By the second day post-infection all insects had succumbed to infection and were dead. Consistent with the initial degradation of the insect cadaver by bacteria predicted by the trophic analysis conducted above, the nematodes began to reproduce by Day 4, 3 days after the insects had died from infection (Fig. 3). At Day 7 post-infection, insect cadavers were placed in a collection trap to encourage the emergence of progeny *S. carpocapsae* IJs. Adults and IJs were observed at this stage. This is when the 2^nd^ generation of nematodes begin to emerge, and endotokia matricida is occurring when some of these juveniles will consume their parents for nutrients. By Day 16 the insect cadavers were largely consumed, and most remaining IJs will have exited. On Day 16 the *S. carpocapsae* IJs, colonized by *X. nematophila* symbionts, were collected from the water trap and analyzed with the other samples. The proportions of nematodes observed at these stages are consistent with previous observations, in which there are more adults and juveniles during the middle phase compared to IJs which changes to high numbers of IJs into the late phase (8). The above observations led us to design a time course metabolomics experiment in which samples were divided into 3 major time frames: the early infection, characterized by bacterial replication and the killing of the insect host, middle infection, characterized by nematode reproduction and nutrient conversion of the cadaver, and late infection, characterized by nutrient depletion and IJ emergence.

**Figure 3:**
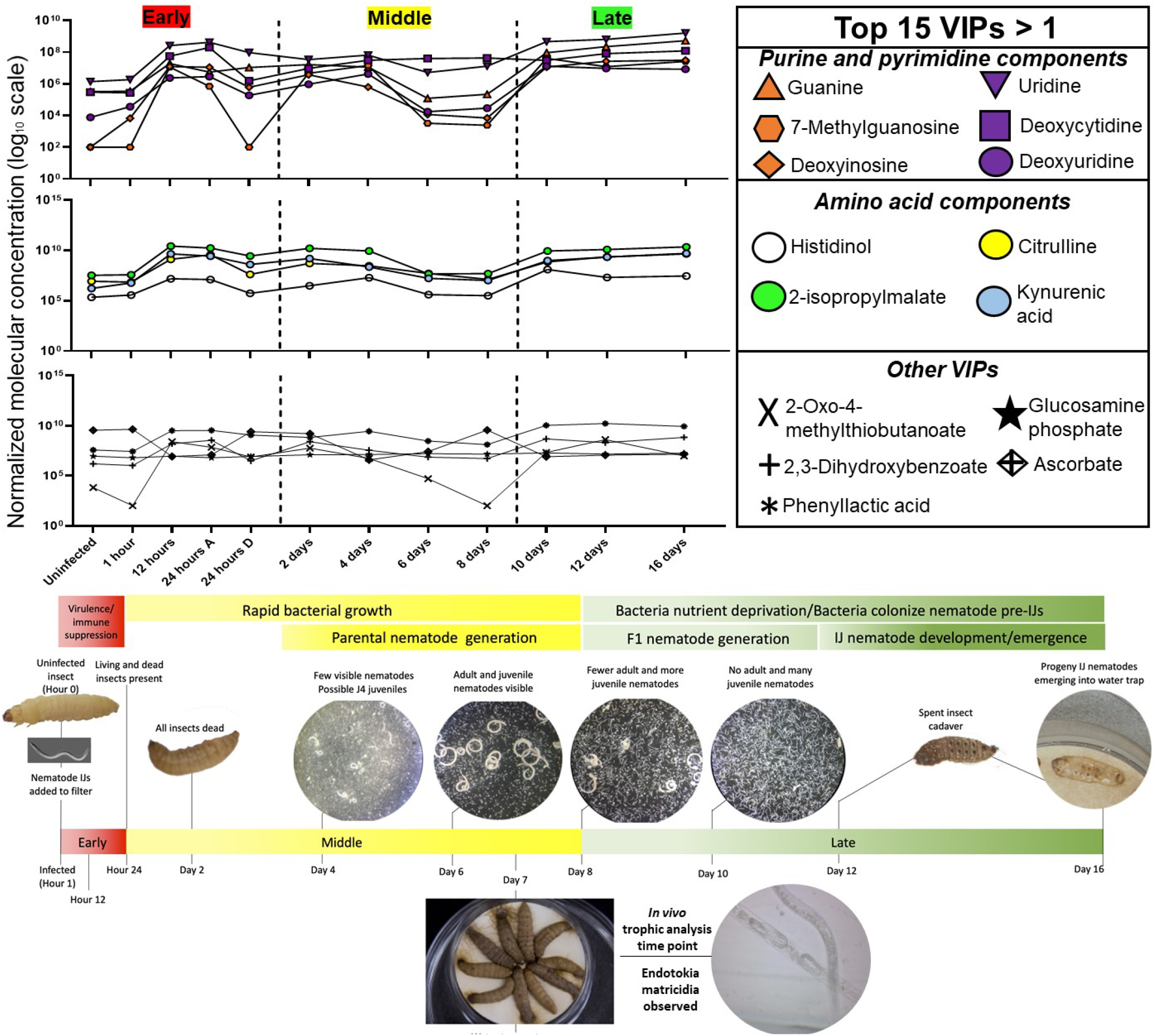
Key moments in the EPNB lifecycle mapped onto important molecules are indicative of the bioconversion of the insect cadaver. The top 15 VIPs>1 metabolites averaged relative abundances were grouped together into 3 categories: purine and pyrimidine components, amino acid components, and other important molecules. Relative metabolite abundance in log scale is displayed on the y-axis of the line graphs.

Metabolites were extracted from individual insects sampled over the infection time course and analyzed using an untargeted UHPLC-HRMS method. The untargeted metabolic profiling analysis revealed 13,748 spectral features. Through the mass spectrometric measurements, a total of 170 of these features were identified based on comparison to known exact mass-to-charge (*m/z*) ratio and retention times from a database of central energy metabolites (Data S3). Another 3,138 unidentified spectral features were included in the analysis and putatively annotated based on their exact masses compared to a *Xenorhabdus* secondary metabolite database (Data S4). This database serves as a rich repository to explore secondary metabolite temporal flux in the tripartite ecosystem of insect, bacteria, and nematode.

### Multivariate data analysis shows metabolic profile gradient corresponding to infection progression

Partial least squares-discriminant analysis (PLS-DA) was performed to observe gross chemical environment changes over time of insect bioconversion to nematode-bacterium complex, when combining all detected metabolite data. A three-dimensional PLS-DA plot shows a progression of distinct metabolic profiles from uninfected insects (black circles) to insects in which bacteria and nematodes are reproducing (red and yellow gradients), and finally to fully consumed insects (green gradients) from which bacterial-colonized infective juvenile populations are emerging (Fig. 4A). Component 1 is 41.1% and contributes the most significantly to the separation observed in the PLS-DA plot.

**Figure 4:**
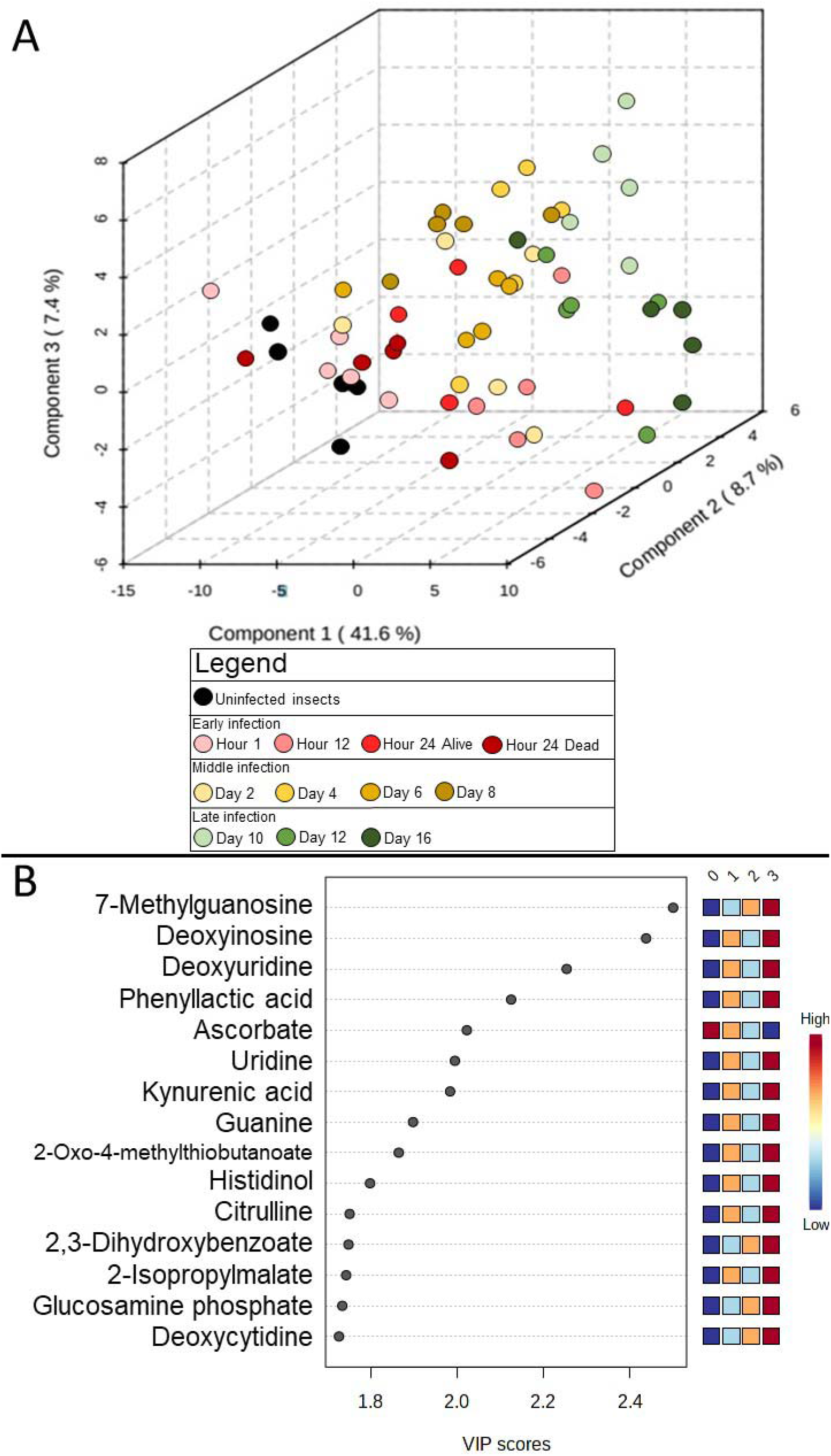
Distinct chemical environments occur during bioconversion of an insect cadaver by *S. carpocapsae* and *X. nematophila*. A) Three-dimensional partial least squares-discriminant analysis (PLS-DA) of time course infection metabolic profiles grouped according to stage of infection: uninfected (black) and early (red gradient), middle (yellow gradient), and late infected insects (green gradient). Components contributing to the separation of the profiles are listed (in %) on the axes. B) The top 15 VIPs contributing to component 1 are listed, where the relative abundance shifts over the time course shown in a heatmap on the right. The numbers on top of the heatmap show the time phases: 0 (uninfected), 1 (early), 2 (middle), and 3 (late).

To examine which metabolites are responsible for most of the variation represented by the PLS plots, Variable Importance in Projection (VIP) values for component 1 were calculated. VIP is a weighted sum of squares of the PLS loadings that considers the amount of explained Y-variation in each dimension. A VIP score >1 indicates that the metabolite significantly contributed to time point differentiation. Most of the VIP>1 metabolites exhibited a bimodal pattern, going from very low in the uninfected insect, to rising in the early bacterial replication phase, to dropping during the middle nematode reproduction phase, and finally rising very high in the late nutrient deplete phase (Fig. 4B and Data S5). Overall, these metabolites were involved in nucleotide and nucleoside biosynthesis, NAD^+^ biosynthesis, and iron acquisition. These included the purine and pyrimidine metabolites 7-methylguanosine, guanine, deoxyinosine, uridine, deoxyuridine, and deoxycytidine. Other top metabolites include kynurenic acid and anthranilate which are precursors to NAD^+^ synthesis. Of the top 15 VIPs, ascorbate was the only molecule to exhibit a decreased abundance over time, dropping from very high abundance to very low later in the time course. This vitamin is necessary for neuron development and could be salvaged from the cadaver to build the nematode nervous systems (34).

Kynurenic acid is an intermediate in the kynurenine pathway and is a way for organisms to synthesize NAD^+^ if they cannot *de novo* synthesize the compound through encoding a quinolinate phosphoribosyltransferase (QPRTase) (35). *S. carpocapsae*, like *C. elegans*, lacks a standard QPRTase but encodes the uridine monophosphate phosphoribosyltransferase (*umps-1*) which synthesizes NAD^+^ from the kynurenine pathway (35, 36). Significant flux of this metabolite throughout the lifecycle could be a metabolic signature of nematode NAD^+^ production.

Phenylacetic acid is a uremic toxin that builds up in kidney patients and is the product of bacterial metabolisms. In *P. aeruginosa* phenylacetic acid (PAA) accumulates at high cell density and inhibits the Type III Secretion System (T3SS), which is toxic to host cells (37). *X. nematophila* does not have a T3SS but does have the evolutionarily related flagellar export apparatus. The transcription factor Lrp positively regulates the flagellar regulon, as well as the gene encoding the XlpA lipase, an enzyme associated with the ability of *X. nematophila* to support *S. carpocapsae* reproduction (19, 20). Although not detected by microarray analysis, quantitative reverse transcriptase analyses indicate that the transcription factor LrhA also positively regulates *xlpA* expression (19). The relatively higher intensity of phenylacetic acid at later stages of insect bioconversion may signal inhibition of secretion of bioconversion enzymes.

2,3-dihydroxybenzoate is a compound involved in siderophore biosynthesis non-ribosomal peptide biosynthesis of siderophore, highlighting a potential importance in iron acquisition from the cadaver. Iron itself does not appear to be limited in the cadaver but it needs to be harvested by the bacteria to aid in nematode reproduction (38). *X. nematophila* does not encode *entA* or *entB*, genes that participate in the conversion of 2,3-dihydroxybenzoate to the enterobactin siderophore pathway (36). Siderophore production is necessary for antibiosis in the closely related *Photorhabdus*-*Heterorhabditis* EPNB symbiosis, which has characterized *phb* genes that are homologous to the *ent* genes (38). The *Photorhabdus phb* genes encode proteins that sequester iron from the cadaver to fend off soil-dwelling bacterial colonizers that exploit the cadaver, and *X. nematophila* does not have any homologs to these genes. *X. nematophila* does not completely dominate the bacterial community of the insect host, suggesting that other taxa such as *Alcaligenes* are possibly responsible for this concentration shift (39).

To identify patterns and groups of metabolite abundance changes over time, hierarchical clustering was performed to reveal groups of metabolites that exhibit similar concentration changes over the time course of bioconversion. A dendrogram of all 170 identified metabolites was generated using the absolute value of the spearman correlation between molecular concentrations, where distance between molecules is defined as 1-|rs| with rs as the spearman rank correlation between time course data points of said molecules (Fig. 5A). Metabolite concentration averages were taken for the four time-phases defined: uninfected, early, middle, and late infection. Metabolite clusters were visualized in a heatmap that displays their pairwise correlation between each molecule (Fig. S2). A heatmap that shows the metabolite clusters, separated by black bars, with the molecule trends in concentration change over time was generated (Fig. 5B). There were 10 total metabolite clusters identified, each with a clear molecular concentration pattern in which the metabolites in that cluster exhibited similar rates of change together over the time phases.

**Figure 5:**
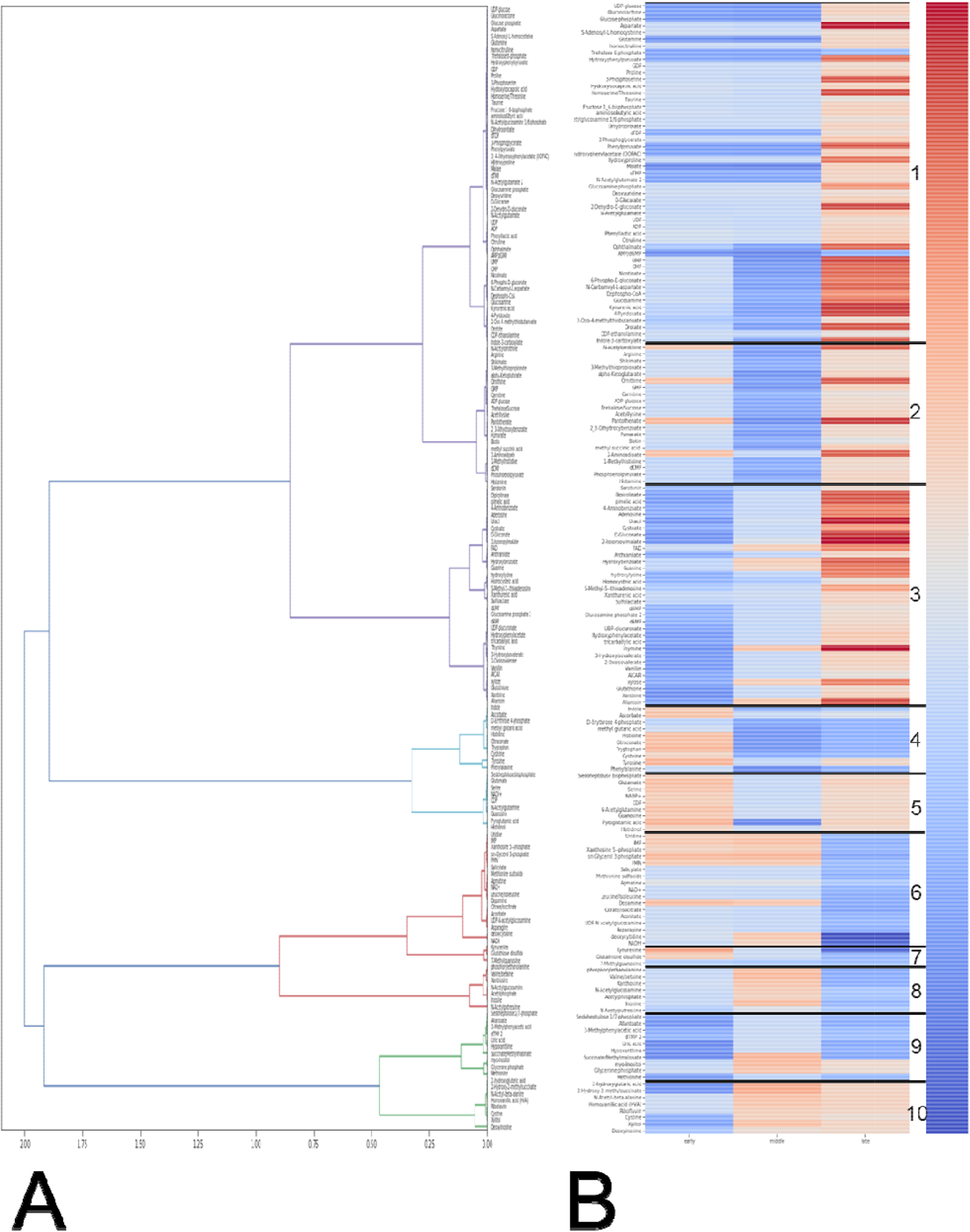
Hierarchical clustering analysis of detected metabolites found ten metabolite cluster exhibit similar rates of change over the infection. A) Dendrogram corresponding to spearman correlation values for each metabolite. B) Identified metabolite clusters with similar log(rate of change) over the lifecycle for the time phase, compared to the previous time phase. Metabolites with a red box exhibit an increased molecular rate of change and metabolites with a blue box exhibit a decreased molecular rate of change.

Clusters of metabolites that exhibit similar rates of change for each time phase were examined to gain an understanding of very broad metabolic pathways affected at each time phase (Data S3). In the early infection phase relative to the previous uninfected phase, there are increased abundances of Clusters 4 and 5. These clusters contain metabolites involved in glutathione biosynthesis (glutamate, cysteine, pyroglutamic acid, and NADP^+^). Glutathione intermediates are increasing while glutathione itself (Cluster 3) is decreasing. Decreased abundance of glutathione paired with increased abundance of synthesis intermediates could be indicative of how the insect is fighting off this pathogen; the insect immune response attempts to strike an equilibrium between non-specific reactive molecules released through the phenoloxidase melanization cascade and the antioxidant glutathione which protects against the reactive molecules (40). Also increasing during this phase are several amino acids, namely those involved in tryptophan metabolism (tryptophan itself and indole). Metabolites that exhibited continuously decreasing abundances in the early phase relative to the uninfected phase were mapped onto Clusters 1, 3, 8, 9, and 10. These clusters contain many compounds, and the highest proportions are involved in purine and pyrimidine biosynthesis and ascorbate metabolism (myo-Inositol, UDP-glucose, UDP-glucuronate, and glucarate). From the microarray data, generally the regulator mutants caused transcripts in ascorbate (*nilR, lrp*) and tryptophan (*lrhA, rpoS*) to decrease. These data from the early phase represent the clusters of metabolites that could be targeted by the bacteria to induce insect death and the metabolites used by the insect to fight the losing war.

From the early phase into the middle infection phase, few clusters exhibit an increase in rates of change. These metabolites are in Clusters 8 and 10, which contain several purine components (deoxyinosine, xanthosine, and inosine) as well as one of the only B vitamins detected in this screen, riboflavin (vitamin B_2_). Other detected B vitamins, like biotin (vitamin B_7_), pantothenate (vitamin B_5_), and 4-pyridoxate (catabolic product of vitamin B_6_), are decreasing in abundance during the middle phase. Many other compounds decreased in abundance during the middle infection phase, relative to the previous early infection phase. These include amino acids (arginine, phenylalanine, tyrosine, tryptophan, cysteine, and methionine), which could reflect that these are the amino acids being incorporated into protein creation for bacterial and nematode biomass accumulation. Microarray analysis indicates widespread differential regulation in these amino acid categories, particularly in the secondary form, *lrp*, and *lrhA* mutants. Other decreasing compounds include pyrimidine intermediates (UMP, CMP, CDP, and UDP) and ascorbate and sugar acid compounds. These data from the middle phase are reflective of the compounds that are being syphoned from the cadaver and incorporated into the nematode lifecycle.

As the insect cadavers entered the late infection phase, nucleic acids (guanine, thymine, and uracil) and amino acids (arginine, cysteine and methionine, and aspartate) steeply increased. This suggests these accumulating compounds are available for nematode DNA, RNA, and protein incorporation, but other factors such as overcrowding in the cadaver or lack of other necessary resources force the nematode to exit. Decreasing rate of change of metabolites relative to the previous middle phase included compounds involved with leucine metabolism and the TCA cycle. These could be more rate-limiting compounds, where their decreasing abundance could signal to the expanding nematode population that it is time to exit.

### Interference of insect tricarboxylic acid (TCA) cycle is critical for infection success and subsequent propagation of nematodes

To identify significant metabolites that are important for infection progression, an ANOVA with post-hoc Tukey’s HSD test was performed on metabolite abundances throughout the lifecycle. Two comparisons were examined: metabolite abundances from uninfected insects compared to individual time points, and individual time points compared to the next subsequent time point. As summarized in Figure 6, TCA cycle components significantly (*p*<0.05) fluctuate in relation to uninfected insects as well as between time phases, throughout the time course. Additionally, significant flux in amino acid metabolism were identified. These trends, especially pertaining to proline and leucine biosynthesis, reveal the importance of insect bioconversion into building blocks essential for nematode development (Supplementary text).

**Figure 6:**
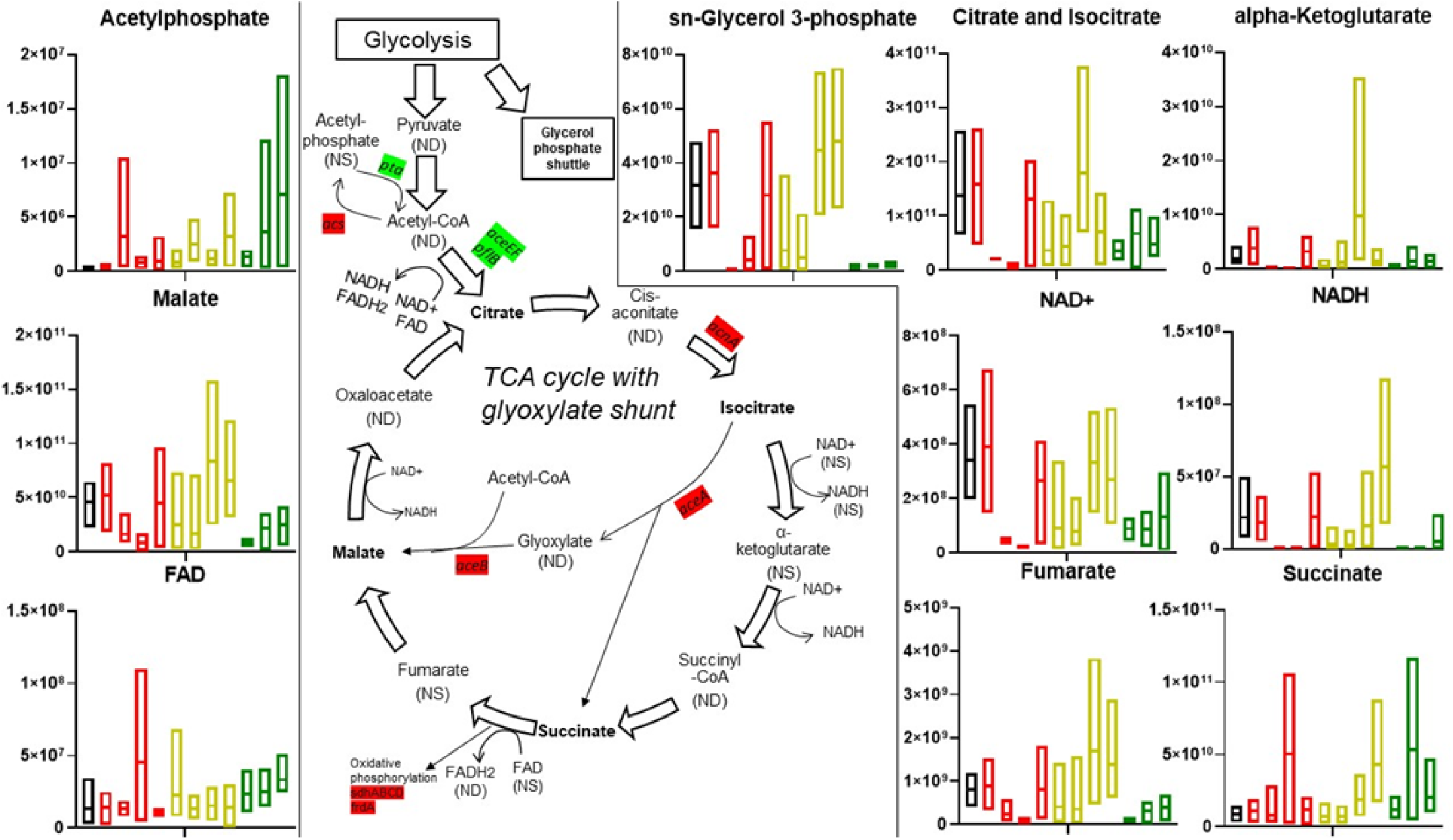
Infection with *S. carpocapsae* IJs affects insect TCA cycle. Normalized molecular concentration box plots throughout the lifecycle are shown for all detected metabolites involved in the TCA cycle. Box plot colors represent which time phase the individual plots belong to for: uninfected (black), early infection (red, going from earliest, 1hr, to latest, 24 hours dead, time points), middle infection (yellow, days 2-8), and late infection (green, days 10-16). Lines in the middle of the boxes indicate the mean molecular concentration. Bolded metabolites indicate significant concentration shifts during the life cycle, as determined by two-way ANOVA (*P*<0.05). NS indicates detection, but not significantly affected (through the ANOVA) throughout infection. ND indicates not detectable. Highlighted genes were detected as significant for the microarray in the Δ*lrhA* and Δ*rpoS* strains. Green indicates positive regulation, red indicates negative regulation.

In the early phase of infection, while the insect is still alive and combatting bacteria and nematode invaders using innate immunity, several key TCA cycle intermediates are reduced in abundance relative to an uninfected insect (Fig. 6). This is shown through significantly decreased abundances of citrate in the Hour 12 and Hour 24 living insects compared to the uninfected insect and the Hour 24 dead insects. Although not significant, a similar trend is observed for two other TCA-related metabolites, malate and *sn*-glycerol-3-phosphate, which aids in NAD^+^ regeneration through the glycerol phosphate shuttle, as well as NAD^+^, NADH, fumarate. This could mean these metabolites were diverted for the immune response, given the differences between the living and dead insects. As the infection progresses into a middle phase, citrate abundances flux, but generally are decreasing. Into the late phase on Day 10, malate, *sn*-glycerol phosphate, succinate, and citrate all drop, which could suggest carbon is being stored (rather than used) in the IJs before they exit the cadaver.

TCA cycle components were mostly in Cluster 6 (*sn*-glycerol-3-phosphate, NAD^+^, NADH, citrate and isocitrate) and Cluster 2 (fumarate, alpha-Ketoglutarate). Student t-tests were utilized to determine additional significant components by comparing each broad time phase to the uninfected insects (Data S3). Acetyl-phosphate was found to be approaching significantly high (*p*<0.1) at the early phase and was significantly high (*p*<0.05) at the middle and late phases. NAD^+^ was found to be approaching significantly low (*p*<0.1) at the early and middle phases and was significantly low (*p*<0.05) for the late phase. *Xenorhabdus* spp. cannot synthesize NAD^+^ and requires nicotinate for growth (41). Generally, the trend for all TCA components in the lifecycle seem to be decreasing as the infection progresses with the exceptions of acetyl-phosphate, FAD, and succinate.

The aforementioned *X. nematophila* mutants experienced differences in transcripts involved in either pyruvate metabolism, glyoxylate metabolism, or the TCA cycle (Data S2). Most of these genes were regulated in the Δ*lrhA* and Δ*rpoS* mutant backgrounds. *aceA* and *aceB* are negatively regulated, while *aceE* and *aceF* were positively regulated between Δ*lrhA* and Δ*rpoS* mutant strains and WT. These genes are involved in the glyoxylate shunt which is a pathway utilized by many bacteria and nematodes to convert 2-carbon compounds into energy resources, in the absence of bountiful sugars (42). Δ*rpoS* mutants upregulate several succinate dehydrogenase genes which are necessary for oxidative phosphorylation. In *E. coli* these genes are regulated in response to different environmental conditions like iron and heme availability (43).

Acetyl-coA is an important node in metabolism, connecting glycolysis, the TCA cycle, fatty acid, amino acid, and secondary metabolite pathways, and acetate dissimilation (44). RpoS and LrhA positively influence the expression of AceF, PflB, Pta, and AckA. Pta-AckA comprise the acetate dissimilation (excretion) pathway (44). Coordinated elevation of these enzymes is predicted to result in lower levels of acetyl-coA and higher levels of acetylphosphate and acetate, which is excreted and available for use by the nematodes. Acetylphosphate is a phosphoryl donor for some response regulators and can be a donor for protein acetylation. Protein acetylation, a ubiquitous post-translational modification in prokaryotes and eukaryotes, is involved in regulation of many different bacterium-host interactions like chemotaxis, replication, and acid resistance, as well as regulating bacterial DNA-binding and protein stability (45). Acetylphosphate was detected in the metabolome and generally increased over the infection, as well as being a VIP>1 metabolite for components 1 and 2. Acetate freely diffuses across membranes and can be incorporated into biomass of both bacteria and nematodes via the glyoxylate shunt (46). *pflB* is predicted to encode the pyruvate-formate lyase (PFL) enzyme involved in conversion of pyruvate and CoA into formate and acetyl-coA and is greatly (>7 |fold change|) downregulated in the Δ*rpoS* and Δ*lrhA* mutants relative to wild-type. PflB converts glucose to formate, and up to one-third of the carbon procured from glucose is converted through this enzyme in *E. coli* (47). PFL condenses acetyl-CoA and formate, allowing for the microbes to use acetate and formate (fermentation products) as the sole carbon sources (48). RpoS and LrhA negatively regulate Acs, which is the acetate assimilation pathway (44). In *E. coli*, Acs activity is inhibited by acetylation of a conserved lysine and its abundance is negatively regulated by the small RNA SdhX (49). RpoS and LrhA also both negatively regulate the TCA cycle enzymes AcnA and AceAB, and this inhibition is predicted to result in accumulation of citrate. Citrate and isocitrate progressively decrease in abundance over the infection cycle and these combined data might indicate that citrate produced and accumulated by *X. nematophila* bacteria may be a provision for nematodes, consumed during reproduction.

## Discussion

A comprehensive framework to understand how metabolism shifts during infection lifecycles of entomopathogenic nematodes was established. Physiologically, it was examined that *X. nematophila* bacteria consume insect tissues, while *S. carpocapsae* nematodes consume bacteria. The high TP_glu-phe_ of 4.5 observed in the nematodes emerging from an insect cadaver suggests the IJs were potentially cannibalizing previous generations of nematodes and/or feeding upon bacteria that were, themselves, already feeding on previous generations of nematodes and bacteria, since if the colonizing nematodes were feeding on bacterial and insect biomass only, they would register at around 3.5. Thus, this high trophic level suggests either endotokia matricida (or bagging) in which nematode eggs hatch within and consume the mother, likely during the second generation of nematodes when nutrients are becoming depleted, or that some other form of cannibalism is occurring within the cadaver (50). A TP_glu-phe_ of 4.6 is similar to many apex predators, such as large marine carnivores or the rare top predators observed in terrestrial ecosystems (51, 52). This underscores the importance of including microbes in studies of organismal trophic identity. In effect, the cadavers used in this study may represent microcosms of the broader communities and ecosystems in which they are embedded. The insect cadavers were, when alive, herbivores. To find multiple levels of carnivory within a single cadaver suggests that a nematode-colonized arthropod mirrors the trophic richness of the broader food-web. The interdigitation of microbial carnivores in a trophic hierarchy—here, nematodes and bacteria—is likely a much more common feature of food-webs than previously thought (28, 53). The microbial trophic identities reported in this study may necessitate a re-calibration of organismal niche concepts, but in so doing, will facilitate the unification of the macro- and microbiome in food web ecology.

It should be emphasized that it has been exceedingly uncommon to find higher-order consumers (TP > 4.0) in a community or ecosystem, given that apex predators feed upon other predators that have, themselves, had to find and subdue ‘lower’ carnivores (54). In classical food web ecology, apex predators are generally considered to be large, fierce, and rare vertebrates (55). However, perhaps the assumption that apex predators exist only within the province of large/fierce/rare vertebrates needs to be revisited. The high trophic positions exhibited by the nematodes in this study suggest that such obligate higher-order consumers are more common than previously thought, with multitudes of apex carnivores existing underfoot in many terrestrial ecosystems. Further, the nematodes can be viewed as farming their symbionts: acting as shepherds that bring their bacterial flock to a fresh insect pasture for harvesting of nutrients.

The trophic study established the foundation to understand metabolic shifts occurring in the cadaver. The time course metabolomics study sought to better understand the process of bioconversion in the cadaver, how is the insect biomass being converted to bacterial and nematode biomass? Applying multivariate statistical tests to the infection metabolomics data set revealed distinct time phase clustering. The variance among the time phases seems to increase as infection progresses, as the healthy insects degrades into bacteria and nematode tissue. Nematode IJ samples were also compared to these time phases. These samples form a distinct cluster away from the other time phases (Fig. S3). The IJ samples completely removed from the lifecycle seem to have metabolic profiles most similar to the late time phase, which are insect samples bursting with nematodes (Fig. 3). However, it is important to note that these are input IJs that have spent weeks outside of an insect cadaver. Reassessing this analysis with output IJs that are recently from the nematode is important to follow up on.

Metabolic analysis revealed TCA cycle components were among the most significant results, indicating that their use by the entomopathogenic nematodes is paramount to infection success and subsequent nematode propagation. Citrate metabolism is ubiquitous in many intracellular pathogens and has been found to be involved in regulating virulence (56). Citrate is necessary for virulence and growth of the Gram-negative pathogenic bacteria *Pseudomonas aeruginosa*, where NADH levels were reduced when the bacterium was treated with citrate and host-killing activity was abolished as a result (57). This group hypothesized that this could be due to decreased flux through the glyoxylate bypass, which has been found to activate the T3SS in this system (58). As mentioned, *X. nematophila* does not encode a T3SS, but does have the evolutionarily related flagellar export apparatus. Several *X. nematophila* glyoxylate bypass genes, were differentially transcribed between avirulent genetic mutants and WT. Glyoxylate was not detected in our screen, and whether flux through this pathway affects virulence should be investigated further. The significant flux of TCA metabolites during the middle phase may be indicative of the role of the TCA cycle in *S. carpocapsae* development, as the TCA cycle has been shown to be essential in early embryogenesis in *C. elegans* (59, 60). Citrate synthase (*cts-1*) and cyclin-dependent kinase 1 (*cdk-1*) were inhibited in these studies and halted *C. elegans* development, and both genes have orthologs in *S. carpocapsae*. Neutral lipids are formed from *sn*-glycerol-3-phosphate and are the major energy reserve in the closely related *S. feltiae* nematodes (61). Fats are stored as lipid droplets in *C. elegans* dauer larvae intestines and serve as a starvation survival mechanism (62). The previously mentioned glyoxylate bypass forms carbohydrates from fatty acids and has been implicated in extending the lifecycle of *C. elegans* (63), highlighting another role of this TCA vs. glyoxylate switching that could be happening later in the life cycle. Any indication that cholesterol is being synthesized from these intermediates can be attributed entirely to the insect’s wheat germ diet, as the *X. nematophila, S. carpocapsae*, or *G. mellonella* cannot synthesize sterols but require them to grow (64).

Additionally, metabolites such as 2,3-dihydroxybenzoate that are not synthesized by *X. nematophila* or *S. carpocapsae* were found to increase over the infection, past insect death. *X. nematophila* does not encode the genes that convert 2,3-dihydroxybenzoate to the siderophore enterobactin, which bind iron to create the ferric enterobactin (FeEnt) complex (65). However, *X. nematophila* encodes FepB, a periplasmic enterobactin binding protein, as well as FepC, FepD, and FepG, which transport the FeEnt into the cell. This increased abundance of this compound suggests other *G. mellonella* microbiome members that survive infection synthesize a compound that is paramount in iron extraction and could contribute to the overall fitness of this symbiosis.

Metabolic analysis also revealed the significance in proline throughout the lifecycle. Insect hemolymph is rich in proline and is used as a main fuel source in some species of flying insects because of its ability to oxidize carbohydrates (66). Proline can be a signal molecule inducing secondary metabolite biosynthesis in *Xenorhabdus* species. Several known virulence factors and antibiotics are regulated via exogenously supplied proline to *Xenorhabdus* cultures (67). *Xenorhabdus* species may have evolved to use proline in insect hemolymph as a preferred amino acid source capable of enhancing the bacterium’s virulence as well as protecting it from various stressors it meets in the insect body. Proline catabolism has also been implicated in promoting stress responses and modulating innate immunity in *C. elegans*, highlighting a possible mechanism by which *S. carpocapsae* survives reactive oxygen species produced by the insect, bacterium, or themselves (68). Enhanced understanding of proline changes over time in the EPNB lifecycle highlights the multiple roles this amino acid is playing for both *Xenorhabdus* virulence and *Steinernema* protection and reproduction. Additional amino acids were found to exhibit similar concentration shifts over the lifecycle via the hierarchical clustering analysis. This machine learning technique can be improved with higher granularity of sample time points, which would strengthen the software developed for this study and allowing it to be used to study more complex and dynamic chemical environments.

We have shown how the parasitic EPNB infection shapes the insect host metabolism. The trophic hierarchy improved how we understand the parasites replicate in the host and highlights the importance of including microbes in these studies. Through rigorous metabolic pathway reconstruction and multivariate statistics, these results suggest each phase of the symbiosis can be characterized by stage-specific chemical signatures. Future targeted metabolomics experiments on EPNB symbioses should be developed to expand the specific trends elucidated by this study. This work adds to a growing scientific foundation on how symbioses, both mutualistic and parasitic, shape the chemical environments they inhabit.

## Materials and Methods

### Conventional nematode and aposymbiotic nematode production

*S. carpocapsae* nematodes (All strain) were propagated through 5^th^ instar larvae of insect *G. mellonella* and using white trp and conventional IJs were collected by trapping in distilled water at stored at room temperature for <1.5 months (69). To generate aposymbiotic IJs, *X. nematophila* Δ*SR1* mutant were grown in Luria Broth (LB) media overnight at 30°C on cell culture wheel and 600μL of overnight bacterial culture were spread onto each of 10mL of lipid agar to grow into confluent lawn at 25°C for 48 hours. Conventional IJs were surface-sterilized, seeded onto Δ*SR1* mutant lawn on lipid agar plates (5000 IJs per 10mL media), and incubated at 25°C for 7 days in dark for nematode reproduction. Aposymbiotic IJs were collected by water-trapping using distilled water and stored at room temperature in dark (69).

### *In vitro* controlled feeding experiment

To collect bacteria sample feeding on terrestrial C3 plants and yeast-based media, *X. nematophila* wild-type or Δ*SR1* mutant bacteria were grown in the dark yeast soy broth (0.5% yeast extract, 3% tryptic soy broth, and 0.5% NaCl) modified from the bacterial growth media from (70) at 30°C on cell culture wheel. Wild-type bacterial overnight cultures (5mL per condition per biological replicate) were collected into microfuge tubes, spun down at >5000RPM, and washed for three times using 1x PBS buffer by resuspending and spinning down the bacterial pellets. Exactly 600μL of Δ*SR1* mutant were spread onto yeast-soy lipid agar plates (0.5% yeast extract, 3% tryptic soy broth, 1.5% agar, 0.2% MgCl_2_, 0.7% corn syrup, 0.4% soy bean oil, supplemented with 40μg cholesterol at per liter of media) and incubated for 48h at 25°C to grow into a confluent lawn. Bacterial lawns were washed off the agar plate using 1x PBS, pelleted and washed as described above. Three individual tubes of bacterial culture were used per strain as three independent biological replicates. To grow nematodes using a controlled diet, approximately 5000 conventional IJs were surface-sterilized and seeded onto bacterial lawn grown on yeast-soy lipid agar plate as described above. Three individual yeast-soy lipid agar plates were used as three biological replicates for each bacterial condition. Three days post seeding IJs, first generation reproductive stage of nematodes (adult males and females) were collected by flooding the bacterial lawn with 1x PBS buffer to resuspend the nematodes. The nematode suspension were collected in the glass cell culture tube and washed for three times by resuspending in 1x PBS buffer. Seven days post seeding the IJs, second generation IJ progenies were collected using distilled water traps and washed in water for three times (settle by gravity and resuspension).

### *In vivo* feeding experiment and sample collection

To prepare insect controls, *G. mellonella* 5^th^ instar larvae were injected with 10uL of either 1x PBS buffer, yeast-soy broth media, or nothing. Three insect larvae were prepared per condition as three biological replicates. To collect nematodes directly fed on *Galleria* insect tissues, *S. carpocapsae* axenic eggs were extracted from adult female nematodes grown on yeast-soy lipid agar plates. Approximately 6000 axenic eggs were seeded to each of the *Galleria-*tissue agar plate (20% (w/v) frozen *G. mellonella* insects cleaned, blended and filtered; 0.5% (w/v) NaCl; and 1.5% (w/v) agar, supplemented with 50mg/L Kanamycin). Mixed-stages of nematodes were collected by flooding the *Galleria-*tissue agar with 1x PBS to resuspend the nematodes, then washed in 1x PBS and water for 3 times to separate nematodes from insect tissue debris. To establish controlled feeding experiments *in vivo* for bacteria and nematodes, *X. nematophila* overnight cultures (in yeast-soy broth) were diluted in 1xPBS buffer, approximately 10^4^ bacterial cells were injected with or without aposymbiotic nematodes (100 IJs per insect). Insect cadaver injected with bacteria only were directly lyophilized and used as insect-bacteria complex controls (see methods below). Insects with bacteria and nematodes co-injection mixture were used to collect IJ progenies by water-trapping, washing (3x in distilled water), and pelleting the IJ samples. Three to five insects were used for each experimental condition as biological replicates.

### Nematode lyophilization and trophic position analysis

Nematodes from *G. mellonella* were collected by placing infected cadavers in modified White traps in which nematodes migrate into distilled water. Trapped nematodes were transferred to Falcon test tubes and allowed settle into a pellet at the bottom of the tube. Nematodes from plate cultivations were harvested by rinsing with sterile distilled water, transferred to Falcon test tubes, and allowed to settle. Samples of nematodes were stored in water at 10°C no longer than a few days until they were lyophilized. For lyophilization, water was decanted off of the sample until only the undisturbed pellet remained at the bottom of the test tube. The top of the test tube was covered with a Kimwipe held in place with a rubber band before lyophilization for >48 h in a Labconco Freezone lyophilizer. During this time, pressures fell below 0.2 millibar, and temperatures reached −50°C. Once the samples had been thoroughly lyophilized, they were removed from tubes using a laboratory spatula that was sterilized with ethanol and dried with kimwipe after every use. Each individual sample was relocated into a sterile 1.5 ml microfuge tube and stored at room temperature for 1-3 months until shipment to Hokkaido, Japan for analysis.

Trophic position (TP_glu-phe_) estimates were generated using the following equation:

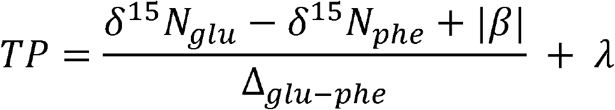

where δ^15^N_glu_ represents the nitrogen isotopic ratio of glutamic acid, δ^15^N_phe_ represents the nitrogen isotopic ratio of phenylalanine, β corrects for the difference in ^15^N values between glutamic acid and phenylalanine within the primary producers of the food web (e.g. β∼8.4‰ for C3 plants), Δ_glu-phe_ represents the net trophic discrimination between glutamic acid and phenylalanine, and λ represents the basal trophic level (=1) of the food web (52). The trophic discrimination factor, Δ_glu-phe_ (referred to here as the TDF_glu-phe_), represents the net intertrophic ^15^N-discrimination between glutamic acid and phenylalanine. Significant differences between known and observed TP values were examined using univariate ANOVA and nonparametric tests (paired Wilcoxon signed rank tests where data were heteroscedastic). Distinguishing among TDF values was accomplished using paired t tests (71).

### Metabolomics sample collection

As per normal infection protocols, 11 *G. mellonella* larvae (Grubco) were placed in the bottom of each of six 6 x 1.5 cm petri plates lined with 2 pieces of #1 filter paper. The filter paper was then inoculated with 1 ml of conventional *S. carpocapsae* IJ stage nematodes (carrying *X. nematophila* bacteria in their intestinal receptacle) to achieve a final average concentration of 10 IJ/μl. At each specified time point (see below), one *G. mellonella* was taken from each of plates 1-5. All insect samples were flash frozen using a dry ice-ethanol bath, and subsequently stored at −80□C. The uninfected, Hour 1 post-infection and Hour 12 post-infection data points were taken of live *G. mellonella*. Since the *G. mellonella* were starting to succumb to the infection at Hour 24, one living and one dead insect was taken at this time point. At Day 7 post-infection, a water trap was set up to enable IJ emergence. At Day 12, the last of the *G. mellonella* from plates 1-5 was used, so insects representing the Day 16 time point were taken entirely from plate 6. Input *S. carpocapsae* IJ and *X. nematophila* symbionts were also collected from the lab stocks and sent for analysis, approximately 50 μl of settled IJ per sample, and a total of 4 samples were sent.

### Preparation for mass spectrometry

For metabolite extraction, *G. mellonella* insects were equilibrated to −20°C for ∼1 h, 300 μl of extraction solution (40:40:20 acetic acid, methanol, and water) was added, and insects were ground using a pestle that fit snugly into the sample tube. All manipulations were performed in a cold room and samples were processed in groups of 12. After grinding, to each tube an additional 1000 μl of extraction solution was added and vortexed for 5-10 sec before being placed at −20°C for 20 min. Tubes were centrifuged at 16,200 x g for 5 min, and the supernatant was decanted to a clean tube. To the original insect sample an additional 200 μl of extraction solution was added, mixed with a pipette tip, vortexed for 5-10 sec, and incubated at −20°C for 20 min. After pelleting the supernatant was combined with the first supernatant sample. Samples were dried (Savant), resuspended, and randomized samples were analyzed consecutively by mass spectrometry using an established 25-minute method (72).

### Metabolomics analysis

An established untargeted metabolomics method utilizing ultra-high performance liquid chromatography coupled to high resolution mass spectrometry (UHPLC-HRMS) (Thermo Scientific, San Jose, CA, USA) was used to analyze water-soluble metabolites (71). A Synergi 2.6 μm Hydro RP column 100 Å, 100 mm x 2.1 mm (Phenomenex, Torrance, CA) and an UltiMate 3000 pump (Thermo Fisher) were used to carry out the chromatographic separations prior to full scan mass analysis by an Exactive Plus Orbitrap MS (Thermo Fisher). HPLC grade solvents (Fisher Scientific, Hampton, NH, USA) were used. Chromatographic peak areas for each detected metabolite were integrated using an open-source software package, Metabolomic Analysis and Visualization Engine (MAVEN) (73, 74). Area under the curve (AUC) was used for further analyses.

### Bacterial strains, plasmids, and culture conditions

Table S1 lists strains used for this study with references where they were originally published. Unless specifically mentioned, *E. coli* were grown in LB broth or on LB plates at 37°C; *X. nematophila* were grown in LB broth or on LB plates supplemented with 0.1% pyruvate at 30°C and kept in dark. Where appropriate, the following antibiotic concentrations were used: ampicillin, 150 μg/ml for *E. coli* and 50 μg/ml for *X. nematophila*; chloramphenicol, 30 μg/ml; erythromycin, 200 μg/ml; kanamycin, 50 μg/ml and streptomycin, 25 μg/ml. *E. coli* donor strain S17 (λpir) or Δ*asd* strain BW29427 was used to conjugate plasmids into *X. nematophila*.

### Microarray experiment and data analysis

Bacteria cultures were grown overnight in 3 ml of LB supplemented with 0.1% pyruvate and appropriate antibiotics in culture tubes at 30°C on roller, subcultured 1:100 into 30 ml of LB supplemented with 0.1% pyruvate and 50 g/ml ampicillin in 125 ml glass flasks and grown for 12 hours to early stationary phase (OD 2-2.1) at 30°C at 150 rpm on shaker. 1 ml of each culture was used to extract total RNA using Qiagen RNeasy Mini Kit, and on-column DNA digestion was performed using Qiagen RNase-Free DNase Set according to manufacturer’s protocol (Qiagen, Valencia, CA). The RNA purity was tested by measuring 260 nm/280 nm and 260 nm/230 nm ratios in TE buffer and the values should be over 1.8. RNA integrity was verified by running 2 g of RNA samples on 1% denaturing agarose gel. The samples were then submitted to Roche NimbleGen for processing and microarray analysis. Gene signals for *lrhA, lrp*, and secondary form *X. nematophila* were compared to HGB800 using a 2-fold change average signal strength cutoff. The *rpoS* mutant was compared to HGB007 using the same significance cutoff. Genes were annotated via the Magnifying Genomes (MaGe) microbial genome annotation system (75), the STRING database (76), as well as through BlastKOALA (33).

### Statistical analysis

PLS-DA plots were generated in MetaboAnalyst 4.0 on August 3rd, 2020. VIP scores were calculated for each component. When more than components are used to calculate the feature importance, the average of the VIP scores are used. The other importance measure is based on the weighted sum of PLS-regression. The weights are a function of the reduction of the sums of squares across the number of PLS components (77). Samples were normalized before processing through MetaboAnalyst based on insect weight. Data was log transformed and pareto scaling was applied. Two-way ANOVA with multiple comparisons and Tukey post-hoc tests were completed by taking individual time point metabolite abundances and comparing their means to the uninfected insect model and each other. Student t-tests were performed by comparing uninfected samples to each time phase (early, middle, and late infection). Relevant metabolic pathways were identified in MetaboAnalyst’s “Pathway Analysis” module using *Drosophila melanogaster, Caenorhabditis elegans*, and *Escherichia coli* as KEGG pathway libraries (78).

## Supporting information

Trophic study results

Microarray results

Time course metabolomics known data

Time course metabolomics unknown data

VIP metabolites derived from the PLS-DA plots

## Acknowledgments

We thank Terra Mauer for her help in insect rearing. We thank Xiaojun Lu for preprocessing the microarray data. We thank Jordan Rogerson for processing the metabolomics samples prior analysis.

## Author contributions

Conceptualization: NCM, MC, MRW, MT, SS, SC, HGB

Methodology: NCM, KAJ, MC, MRW, SF, SK, MT, YC, SS, SC

Investigation: NCM, KAJ, MC, MRW, SS, SC, HGB

Visualization: NCM, KAJ, MRW, SS, HGB

Supervision: SK, MT, YC, SS, SC, HGB

Writing—original draft: NCM, SS, HGB

Writing—review & editing: NCM, KAJ, MC, MRW, SF, SK, MT, SS, SC, HGB

## Data and materials availability

All data are available in the main text or the supplementary materials. Additional detail on the hierarchical clustering analysis can be found at: http://doi.org/10.5281/zenodo.3962081

## References

1. L. Margulis, Symbiosis in cell evolution: life and its environment on the early Earth. (W. H. Freeman, San Francisco, 1981), pp. xxii, 419 p.

2. A. Camilli, B. L. Bassler, Bacterial small-molecule signaling pathways. Science 311, 1113–1116 (2006).

3. R. J. Worthington, J. J. Richards, C. Melander, Small molecule control of bacterial biofilms. Org Biomol Chem 10, 7457–7474 (2012).

4. J. Chaston, A. E. Douglas, Making the most of ‘omics’ for symbiosis research. Biol Bull 223, 21–29 (2012).

5. S. E. Maddocks, P. C. F. Oyston, Structure and function of the LysR-type transcriptional regulator (LTTR) family proteins. Microbiology (Reading) 154, 3609–3623 (2008).

6. J. A. Budnick, L. M. Sheehan, M. J. Ginder, K. C. Failor, J. M. Perkowski, J. F. Pinto, K. A. Kohl, L. Kang, P. Michalak, L. Luo, J. E. Heindl, C. C. Caswell, A central role for the transcriptional regulator VtlR in small RNA-mediated gene regulation in Agrobacterium tumefaciens. Sci Rep 10, 14968 (2020).

7. H. Abdelhamed, R. Ramachandran, L. Narayanan, O. Ozdemir, A. Cooper, A. K. Olivier, A. Karsi, M. L. Lawrence, Contributions of a LysR Transcriptional Regulator to Listeria monocytogenes Virulence and Identification of Its Regulons. J Bacteriol 202, (2020).

8. M. Cao, H. Goodrich-Blair, Xenorhabdus nematophila bacteria shift from mutualistic to virulent Lrp-dependent phenotypes within the receptacles of Steinernema carpocapsae insect-infective stage nematodes. Environ Microbiol 22, 5433–5449 (2020).

9. J. Tang, Microbial metabolomics. Curr Genomics 12, 391–403 (2011).

10. V. V. Phelan, W. J. Moree, J. Aguilar, D. S. Cornett, A. Koumoutsi, S. M. Noble, K. Pogliano, C. A. Guerrero, P. C. Dorrestein, Impact of a transposon insertion in phzF2 on the specialized metabolite production and interkingdom interactions of Pseudomonas aeruginosa. J Bacteriol 196, 1683–1693 (2014).

11. Y. Dong, G. J. Brewer, Global Metabolic Shifts in Age and Alzheimer’s Disease Mouse Brains Pivot at NAD+/NADH Redox Sites. J Alzheimers Dis 71, 119–140 (2019).

12. G. R. Richards, H. Goodrich-Blair, Masters of conquest and pillage: Xenorhabdus nematophila global regulators control transitions from virulence to nutrient acquisition. Cell Microbiol 11, 1025–1033 (2009).

13. S. P. Stock, Partners in crime: symbiont-assisted resource acquisition in Steinernema entomopathogenic nematodes. Curr Opin Insect Sci 32, 22–27 (2019).

14. M. Javal, J. S. Terblanche, D. E. Conlong, A. P. Malan, First Screening of Entomopathogenic Nematodes and Fungus as Biocontrol Agents against an Emerging Pest of Sugarcane, Cacosceles newmannii (Coleoptera: Cerambycidae). Insects 10, (2019).

15. W. J. da Silva, H. L. Pilz-Junior, R. Heermann, O. S. da Silva, The great potential of entomopathogenic bacteria Xenorhabdus and Photorhabdus for mosquito control: a review. Parasit Vectors 13, 376 (2020).

16. N. Killiny, Generous hosts: Why the larvae of greater wax moth, Galleria mellonella is a perfect infectious host model? Virulence 9, 860–865 (2018).

17. E. E. Herbert, H. Goodrich-Blair, Friend and foe: the two faces of Xenorhabdus nematophila. Nat Rev Microbiol 5, 634–646 (2007).

18. D. K. Mitani, H. K. Kaya, H. Goodrich-Blair, Comparative study of the entomopathogenic nematode, Steinernema carpocapsae, reared on mutant and wild-type Xenorhabdus nematophila. Biological Control 29, 382–391 (2004).

19. G. R. Richards, E. E. Herbert, Y. Park, H. Goodrich-Blair, Xenorhabdus nematophila lrhA is necessary for motility, lipase activity, toxin expression, and virulence in Manduca sexta insects. J Bacteriol 190, 4870–4879 (2008).

20. G. R. Richards, H. Goodrich-Blair, Examination of Xenorhabdus nematophila lipases in pathogenic and mutualistic host interactions reveals a role for xlpA in nematode progeny production. Appl Environ Microbiol 76, 221–229 (2010).

21. E. I. Vivas, H. Goodrich-Blair, Xenorhabdus nematophilus as a model for host-bacterium interactions: rpoS is necessary for mutualism with nematodes. J Bacteriol 183, 4687–4693 (2001).

22. E. E. Herbert Tran, H. Goodrich-Blair, CpxRA Contributes to Xenorhabdus nematophila Virulence through Regulation of lrhA and Modulation of Insect Immunity. Applied and Environmental Microbiology 75, 3998 (2009).

23. K. N. Cowles, C. E. Cowles, G. R. Richards, E. C. Martens, H. Goodrich-Blair, The global regulator Lrp contributes to mutualism, pathogenesis and phenotypic variation in the bacterium Xenorhabdus nematophila. Cell Microbiol 9, 1311–1323 (2007).

24. C. E. Cowles, H. Goodrich-Blair, nilR is necessary for co-ordinate repression of Xenorhabdus nematophila mutualism genes. Molecular microbiology 62, 760–771 (2006).

25. D. Park, S. Forst, Co-regulation of motility, exoenzyme and antibiotic production by the EnvZ-OmpR-FlhDC-FliA pathway in Xenorhabdus nematophila. Mol Microbiol 61, 1397–1412 (2006).

26. A. Givaudan, S. Baghdiguian, A. Lanois, N. Boemare, Swarming and Swimming Changes Concomitant with Phase Variation in Xenorhabdus nematophilus. Appl Environ Microbiol 61, 1408–1413 (1995).

27. M. C. Cambon, N. Parthuisot, S. Pagès, A. Lanois, A. Givaudan, J.-B. Ferdy, Selection of Bacterial Mutants in Late Infections: When Vector Transmission Trades Off against Growth Advantage in Stationary Phase. mBio 10, e01437–01419 (2019).

28. S. A. Steffan, Y. Chikaraishi, P. S. Dharampal, J. N. Pauli, C. Guedot, N. Ohkouchi, Unpacking brown foodwebs: Animal trophic identity reflects rampant microbivory. Ecol Evol 7, 3532–3541 (2017).

29. C. E. Cowles, H. Goodrich-Blair, The Xenorhabdus nematophila nilABC genes confer the ability of Xenorhabdus spp. to colonize Steinernema carpocapsae nematodes. J Bacteriol 190, 4121–4128 (2008).

30. E. A. Hussa, A. M. Casanova-Torres, H. Goodrich-Blair, The Global Transcription Factor Lrp Controls Virulence Modulation in Xenorhabdus nematophila. J Bacteriol 197, 3015–3025 (2015).

31. X. Lu, “Examining Pathogenic and Mutualistic Regulatory Networks in the Bacterium Xenorhabdus nematophila,” Ph.D. thesis, UW-Madison, Madison, WI (2012).

32. X. Lu, H. Goodrich-Blair, B. Tjaden, Assessing computational tools for the discovery of small RNA genes in bacteria. RNA 17, 1635–1647 (2011).

33. M. Kanehisa, Y. Sato, K. Morishima, BlastKOALA and GhostKOALA: KEGG Tools for Functional Characterization of Genome and Metagenome Sequences. J Mol Biol 428, 726–731 (2016).

34. J. M. May, Z. C. Qu, M. E. Meredith, Mechanisms of ascorbic acid stimulation of norepinephrine synthesis in neuronal cells. Biochem Biophys Res Commun 426, 148–152 (2012).

35. Y. Lee, H. Jeong, K. H. Park, K. W. Kim, Effects of NAD(+) in Caenorhabditis elegans Models of Neuronal Damage. Biomolecules 10, (2020).

36. A. Rougon-Cardoso, M. Flores-Ponce, H. E. Ramos-Aboites, C. E. Martinez-Guerrero, Y. J. Hao, L. Cunha, J. A. Rodriguez-Martinez, C. Ovando-Vazquez, J. R. Bermudez-Barrientos, C. Abreu-Goodger, N. Chavarria-Hernandez, N. Simoes, R. Montiel, The genome, transcriptome, and proteome of the nematode Steinernema carpocapsae: evolutionary signatures of a pathogenic lifestyle. Sci Rep 6, 37536 (2016).

37. J. Wang, Y.-H. Dong, T. Zhou, X. Liu, Y. Deng, C. Wang, J. Lee, L.-H. Zhang, Pseudomonas aeruginosa Cytotoxicity Is Attenuated at High Cell Density and Associated with the Accumulation of Phenylacetic Acid. PloS one 8, e60187 (2013).

38. T. A. Ciche, M. Blackburn, J. R. Carney, J. C. Ensign, Photobactin: a catechol siderophore produced by Photorhabdus luminescens, an entomopathogen mutually associated with Heterorhabditis bacteriophora NC1 nematodes. Appl Environ Microbiol 69, 4706–4713 (2003).

39. M. C. Cambon, P. Lafont, M. Frayssinet, A. Lanois, J. C. Ogier, S. Pages, N. Parthuisot, J. B. Ferdy, S. Gaudriault, Bacterial community profile after the lethal infection of Steinernema-Xenorhabdus pairs into soil-reared Tenebrio molitor larvae. FEMS Microbiol Ecol 96, (2020).

40. K. D. Clark, Z. Lu, M. R. Strand, Regulation of melanization by glutathione in the moth Pseudoplusia includens. Insect Biochem Mol Biol 40, 460–467 (2010).

41. E. C. Martens, F. M. Russell, H. Goodrich-Blair, Analysis of Xenorhabdus nematophila metabolic mutants yields insight into stages of Steinernema carpocapsae nematode intestinal colonization. Mol Microbiol 58, 28–45 (2005).

42. S. R. Maloy, W. D. Nunn, Genetic regulation of the glyoxylate shunt in Escherichia coli K-12. J Bacteriol 149, 173–180 (1982).

43. S. J. Park, C. P. Tseng, R. P. Gunsalus, Regulation of succinate dehydrogenase (sdhCDAB) operon expression in Escherichia coli in response to carbon supply and anaerobiosis: role of ArcA and Fnr. Mol Microbiol 15, 473–482 (1995).

44. A. J. Wolfe, Glycolysis for Microbiome Generation. Microbiol Spectr 3, (2015).

45. J. Ren, Y. Sang, J. Lu, Y. F. Yao, Protein Acetylation and Its Role in Bacterial Virulence. Trends Microbiol 25, 768–779 (2017).

46. J. T. Alaimo, S. J. Davis, S. S. Song, C. R. Burnette, M. Grotewiel, K. L. Shelton, J. T. Pierce-Shimomura, A. G. Davies, J. C. Bettinger, Ethanol metabolism and osmolarity modify behavioral responses to ethanol in C. elegans. Alcohol Clin Exp Res 36, 1840–1850 (2012).

47. C. Doberenz, M. Zorn, D. Falke, D. Nannemann, D. Hunger, L. Beyer, C. H. Ihling, J. Meiler, A. Sinz, R. G. Sawers, Pyruvate formate-lyase interacts directly with the formate channel FocA to regulate formate translocation. J Mol Biol 426, 2827–2839 (2014).

48. L. Zelcbuch, S. N. Lindner, Y. Zegman, I. Vainberg Slutskin, N. Antonovsky, S. Gleizer, R. Milo, A. Bar-Even, Pyruvate Formate-Lyase Enables Efficient Growth of Escherichia coli on Acetate and Formate. Biochemistry 55, 2423–2426 (2016).

49. F. De Mets, L. Van Melderen, S. Gottesman, Regulation of acetate metabolism and coordination with the TCA cycle via a processed small RNA. Proc Natl Acad Sci U S A 116, 1043–1052 (2019).

50. Y. Baliadi, T. Yoshiga, E. Kondo, Infectivity and post-infection development of infective juveniles originating via endotokia matricida in entomopathogenic nematodes. Applied Entomology and Zoology 39, 61–69 (2004).

51. Y. Chikaraishi, N. O. Ogawa, H. Doi, N. Ohkouchi, 15N/14N ratios of amino acids as a tool for studying terrestrial food webs: a case study of terrestrial insects (bees, wasps, and hornets). Ecological Research 26, 835–844 (2011).

52. Y. Chikaraishi, S. A. Steffan, N. O. Ogawa, N. F. Ishikawa, Y. Sasaki, M. Tsuchiya, N. Ohkouchi, High-resolution food webs based on nitrogen isotopic composition of amino acids. Ecol Evol 4, 2423–2449 (2014).

53. S. A. Steffan, P. S. Dharampal, Undead food-webs: Integrating microbes into the food-chain. Food Webs 18, e00111 (2019).

54. R. L. Lindeman, The trophic-dynamic aspect of ecology. Bulletin of Mathematical Biology 53, 167–191 (1942)

55. P. A. Colinvaux, B. D. Barnett, Lindeman and the Ecological Efficiency of Wolves. The American Naturalist 114, 707–718 (1979).

56. G. P. Martino, C. E. Perez, C. Magni, V. S. Blancato, Implications of the expression of Enterococcus faecalis citrate fermentation genes during infection. PLoS One 13, e0205787 (2018).

57. K. Perinbam, J. V. Chacko, A. Kannan, M. A. Digman, A. Siryaporn, A Shift in Central Metabolism Accompanies Virulence Activation in Pseudomonas aeruginosa. mBio 11, (2020).

58. J. C. Chung, O. Rzhepishevska, M. Ramstedt, M. Welch, Type III secretion system expression in oxygen-limited Pseudomonas aeruginosa cultures is stimulated by isocitrate lyase activity. Open Biol 3, 120131 (2013).

59. K. Hada, K. Hirota, A. Inanobe, K. Kako, M. Miyata, S. Araoi, M. Matsumoto, R. Ohta, M. Arisawa, H. Daitoku, T. Hanada, A. Fukamizu, Tricarboxylic acid cycle activity suppresses acetylation of mitochondrial Harris proteins during early embryonic development in Caenorhabditis elegans. J Biol Chem 294, 3091–3099 (2019).

60. M. M. Rahman, S. Rosu, D. Joseph-Strauss, O. Cohen-Fix, Down-regulation of tricarboxylic acid (TCA) cycle genes blocks progression through the first mitotic division in Caenorhabditis elegans embryos. Proc Natl Acad Sci U S A 111, 2602–2607 (2014).

61. D. J. Wright, P. S. Grewal, M. Stolinski, Relative importance of neutral lipids and glycogen as energy stores in dauer larvae of two entomopathogenic nematodes, Steinernema carpocapsae and Steinernema feltiae. Comp Biochem Physiol B Biochem Mol Biol 118, 269–273 (1997).

62. H. Y. Mak, Lipid droplets as fat storage organelles in Caenorhabditis elegans: Thematic Review Series: Lipid Droplet Synthesis and Metabolism: from Yeast to Man. J Lipid Res 53, 28–33 (2012).

63. C. B. Edwards, N. Copes, A. G. Brito, J. Canfield, P. C. Bradshaw, Malate and fumarate extend lifespan in Caenorhabditis elegans. PLoS One 8, e58345 (2013).

64. X. Jing, S. T. Behmer, Insect Sterol Nutrition: Physiological Mechanisms, Ecology, and Applications. Annu Rev Entomol 65, 251–271 (2020).

65. K. N. Raymond, E. A. Dertz, S. S. Kim, Enterobactin: an archetype for microbial iron transport. Proc Natl Acad Sci U S A 100, 3584–3588 (2003).

66. L. Teulier, J. M. Weber, J. Crevier, C. A. Darveau, Proline as a fuel for insect flight: enhancing carbohydrate oxidation in hymenopterans. Proc Biol Sci 283, (2016).

67. J. M. Crawford, R. Kontnik, J. Clardy, Regulating alternative lifestyles in entomopathogenic bacteria. Curr Biol 20, 69–74 (2010).

68. H. Tang, S. Pang, Proline Catabolism Modulates Innate Immunity in Caenorhabditis elegans. Cell Rep 17, 2837–2844 (2016).

69. J. G. McMullen, 2nd, S. P. Stock, In vivo and in vitro rearing of entomopathogenic nematodes (Steinernematidae and Heterorhabditidae). J Vis Exp, 52096 (2014).

70. E. J. Buecher, I. Popiel, Liquid Culture of the Entomogenous Nematode Steinernema feltiae with Its Bacterial Symbiont. J Nematol 21, 500–504 (1989).

71. S. A. Steffan, Y. Chikaraishi, C. R. Currie, H. Horn, H. R. Gaines-Day, J. N. Pauli, J. E. Zalapa, N. Ohkouchi, Microbes are trophic analogs of animals. Proc Natl Acad Sci U S A 112, 15119–15124 (2015).

72. W. Lu, M. F. Clasquin, E. Melamud, D. Amador-Noguez, A. A. Caudy, J. D. Rabinowitz, Metabolomic analysis via reversed-phase ion-pairing liquid chromatography coupled to a stand alone orbitrap mass spectrometer. Anal Chem 82, 3212–3221 (2010).

73. M. F. Clasquin, E. Melamud, J. D. Rabinowitz, LC-MS data processing with MAVEN: a metabolomic analysis and visualization engine. Curr Protoc Bioinformatics Chapter 14, Unit14 11 (2012).

74. E. Melamud, L. Vastag, J. D. Rabinowitz, Metabolomic analysis and visualization engine for LC-MS data. Anal Chem 82, 9818–9826 (2010).

75. D. Vallenet, L. Labarre, Z. Rouy, V. Barbe, S. Bocs, S. Cruveiller, A. Lajus, G. Pascal, C. Scarpelli, C. Medigue, MaGe: a microbial genome annotation system supported by synteny results. Nucleic Acids Res 34, 53–65 (2006).

76. D. Szklarczyk, A. L. Gable, D. Lyon, A. Junge, S. Wyder, J. Huerta-Cepas, M. Simonovic, N. T. Doncheva, J. H. Morris, P. Bork, L. J. Jensen, C. V. Mering, STRING v11: protein-protein association networks with increased coverage, supporting functional discovery in genome-wide experimental datasets. Nucleic Acids Res 47, D607–D613 (2019).

77. J. Chong, J. Xia, Using MetaboAnalyst 4.0 for Metabolomics Data Analysis, Interpretation, and Integration with Other Omics Data. Methods Mol Biol 2104, 337–360 (2020).

78. M. Kanehisa, M. Furumichi, M. Tanabe, Y. Sato, K. Morishima, KEGG: new perspectives on genomes, pathways, diseases and drugs. Nucleic Acids Res 45, D353–D361 (2017).

## Supplementary References

79. Y. Chikaraishi, Y. Kashiyama, N. O. Ogawa, Y. Kashiyama, H. Kitazato, N. Ohkouchi, Metabolic control of nitrogen isotope composition of amino acids in macroalgae and gastropods: implications for aquatic food web studies. Marine Ecology Progress Series 342, 85–90 (2007).

80. Y. Chikaraishi, N. Ogawa, Y. Kashiyama, Y. Takano, H. Suga, A. Tomitani, H. Miyashita, H. Kitazato, N. Ohkouchi, Determination of aquatic food-web structure based on compound-specific nitrogen isotopic composition of amino acids: Trophic level estimation by amino acid δ 15 N. Limnology and oceanography 7, 740–750 (2009).

81. S. A. Steffan, Y. Chikaraishi, D. R. Horton, N. Ohkouchi, M. E. Singleton, E. Miliczky, D. B. Hogg, V. P. Jones, Trophic hierarchies illuminated via amino acid isotopic analysis. PLoS One 8, e76152 (2013).

82. M. Watford, Glutamine metabolism and function in relation to proline synthesis and the safety of glutamine and proline supplementation. J Nutr 138, 2003S–2007S (2008).

83. R. Zheng, X. Feng, X. Wei, X. Pan, C. Liu, R. Song, Y. Jin, F. Bai, S. Jin, W. Wu, Z. Cheng, PutA Is Required for Virulence and Regulated by PruR in Pseudomonas aeruginosa. Front Microbiol 9, 548 (2018).

84. M. L. Di Martino, R. Campilongo, M. Casalino, G. Micheli, B. Colonna, G. Prosseda, Polyamines: emerging players in bacteria-host interactions. Int J Med Microbiol 303, 484–491 (2013).

85. P. Shah, E. Swiatlo, A multifaceted role for polyamines in bacterial pathogens. Mol Microbiol 68, 4–16 (2008).

86. Zecic, I. Dhondt, B. P. Braeckman, The nutritional requirements of Caenorhabditis elegans. Genes Nutr 14, 15 (2019).

