## Supplemental text and figures for "Chemical ecology of an apex predator life cycle"

**This PDF file includes:**

Supplementary Text

Figs. S1 to S2

Tables S1 to S2

References (79 to 86)

**Other Supplementary Materials for this manuscript include the following:**

Data S1 to S5

Supplementary Text

***In vitro* trophic analysis trials**

This past work relied upon compound-specific isotopic analyses of select amino acid pools, particularly the degree of ^15^N-enrichment between two amino acids—glutamic acid (glu) and phenylalanine (phe). The differential enrichment between these two amino acids provides a measure of inter-trophic enrichment, which is largely attributable to an organism’s assimilation of dietary amino acids (81, 82). Such inter-trophic enrichment has been referred to as the trophic discrimination factor (TDF_glu-phe_), and in carefully controlled feeding studies among diverse consumer groups in the Animalia, Fungi, and Bacteria, the TDF_glu-phe_ has averaged approximately 7.2‰ (28, 71). Here, following controlled-feeding *in vitro* trials, the nematodes and bacteria were shown to have both registered TDF_glu-phe_ values in line with past findings (Fig. 1A). Specifically, the mean (± SE) TDF_glu-phe_ value exhibited by nematodes cultured on bacterial lawns was 7.41 ± 0.22‰ (*N* = 14). When parsed by nematode stage, the TDF_glu-phe_ values of adult and infective juvenile nematodes were, respectively, 6.96 ± 0.16‰ (*N* = 8) and 8.02 ± 0.36‰ (*N* = 6). Nematodes fed exclusively a diet of homogenized insect biomass produced a TDF of 7.38 ± 0.05‰ (*N* = 3). Collectively, the nematode TDF was 7.40 ± 0.18‰ (*N* = 17). The bacterial symbiont, *Xenorhabdus*, which had been cultured on agar growth media, registered a TDF_glu-phe_ of 6.53 ± 0.20‰ (*N* = 6). The mean TDF_glu-phe_ across both the nematodes and bacteria in this food-chain was 7.18 ± 0.16‰, which did not represent a significant departure from the generalized 7.2‰ TDF_glu-phe_ benchmark (*t_22_* = -0.14, *P* = 0.893). Given the degree of inter-trophic enrichment exhibited in the nematodes and bacteria, these consumer groups were consistent with the enrichment patterns of heterotrophs across terrestrial, marine, and freshwater systems, allowing for trophic position estimation using established isotopic protocols (28, 51, 71 81-83).

Using compound-specific isotopic analysis of amino acids, the trophic identities of consumers and their respective diets within the *in vitro* food-chain were measured. At the base of the food-chain, the agar growth media registered a trophic position (TP_glu-phe_) of 1.0 ± 0.04‰ (*N* = 3), and the bacteria feeding upon the agar registered at 1.9 ± 0.03‰ (*N* = 6), which represented approximately one trophic level higher than their diet. Correspondingly, the adult and infective juvenile nematodes that had fed upon the bacteria registered, respectively, at 2.90 ± 0.02‰ (*N* = 8) and 3.0 ± 0.06‰ (*N* = 6), which, as predicted, was exactly one trophic level higher than their diet. The homogenate of insect biomass was measured at 2.2 ± 0.02‰ (*N* = 6), and the nematodes feeding exclusively on this homogenate registered at 3.2 ± 0.01‰ (*N* = 4), which again demonstrated that when the nematodes consumed a given diet, they registered one trophic level higher. The *in vitro* food-chain effectively compartmentalized each consumer group and thereby provided a means to confirm that when the nematodes or bacteria consumed a given diet, their isotopic compositions enriched consistently and produced predictable trophic position estimates.

**Analysis of amino acid abundances found proline fluctuates throughout the infection**

The amino acid pyroglutamic acid, which is a precursor to glutamate, significantly rises (as determined through the ANOVA tests) in abundance during the late phase, at Day 16 compared to the uninfected insect and between Day 12 and Day 16. Glutamate is a precursor for proline metabolism (84). Proline abundances fluctuated significantly throughout the time course. Compared to uninfected insects, proline levels were significantly lower at Day 10 and higher at Day 16, but otherwise were not significantly different. When comparing each time point to the previous, there was a significantly higher level of proline in dead insects at Hour 24 relative to living insects at the same time point. However, in dead insects between Hour 24 and Day 2 the levels of proline dropped again. Thereafter there was a cycle of increase and decrease in proline abundance (Day 4<Day 6, Day 8>Day 10, Day 12<Day 16), with an overall rise throughout the middle and late phases. Additional metabolites in the arginine/proline/polyamine metabolism pathways that were significantly different among samples based on student t-tests (Data S3), included ornithine, which was significantly high at the early and middle infection stages compared to uninfected insects, and hydroxyproline, which was significantly high at early, middle, and late infection stages compared to uninfected insects. Hierarchical clustering analysis reveals Cluster 1 (proline and hydroxyproline) and Cluster 5 (glutamate and pyroglutamic acid) contain these compounds. These cluster trends differ in the early phase, where Cluster 1 decreases while Cluster 5 increases, possibly indicating that these compounds are getting converted into each other.

Microarray analysis of Δ*lrhA*, Δ*lrp*, Δ*nilR*, Δ*rpoS*, and secondary form *X. nematophila* mutants reveal similarities to each other in disrupted proline gene regulation compared to their wild type/primary form counterparts. *putA*, predicted to encode 1-pyrroline-5-carboxylate dehydrogenase, an enzyme involved in the conversion of proline to glutamate, was negatively regulated (2<|fold change|) in the *ΔlrhA* and *ΔrpoS* strains, and positively regulated in the secondary form strain. PutA is necessary for virulence in *Pseudomonas aeruginosa* and protects the bacterium against oxidative stress (85). XNC1_2468 is downregulated in the Δ*lrp*, Δ*rpoS*, and secondary form mutant backgrounds. This gene encodes a spermidine N1-acetyltransferase as part of polyamine biosynthesis from ornithine and putrescine. XNC1_2274 and XNC1_3619 are FAD-dependent oxidoreductases involved downstream of proline metabolism in putrescine utilization and are upregulated in the Δ*lrhA*, Δ*lrp*, Δ*nilR*, and Δ*rpoS* mutant backgrounds. These genes are mostly involved in downstream proline metabolism, specifically polyamine biosynthesis. Pathogenic gram-negative bacteria exploit polyamine-related processes of their host for growth and proliferation, using these host molecules for toxin activity, biofilm production, and limiting host immune responses (86, 87). XNC1_2154 is predicted to encode an enzyme with L-aspartate:2-oxoglutarate aminotransferase activity and is downregulated in the secondary form and Δ*lrp* mutant backgrounds.

Amino acids leucine, isoleucine, and phenylalanine amino acids were significantly higher in abundance at the late stage of infection. These are essential amino acids which *C. elegans*, and presumably *S. carpocapsae*, must acquire from its diet (i.e. bacteria) (88). Several *X. nematophila* genes, including *ilvC* and *ilvI*, involved in leucine/isoleucine biosynthesis were differentially regulated in the Δ*lrhA* (negatively) and Δ*rpoS* (positively regulates *ilvC* and negatively regulates *ilvI*) mutant backgrounds, relative to wild type. Δ*rpoS* mutants had higher levels of *leuA* transcript, a gene predicted to encode a 2-isopropylmalate synthase, a leucine precursor which is increases throughout the lifecycle. Δ*lrhA* increased transcript abundances of *fadA, fadB*, *fadI,* and *fadJ* which participate in the conversion of leucine and isoleucine into fatty acids and acetyl-CoA. In the Δ*lrp* and Δ*rpoS* mutant backgrounds, *mmsA* was upregulated, another gene that encodes a methylmalonate-semialdehyde dehydrogenase that participates in the breakdown of leucine and isoleucine.

Fig. S1.

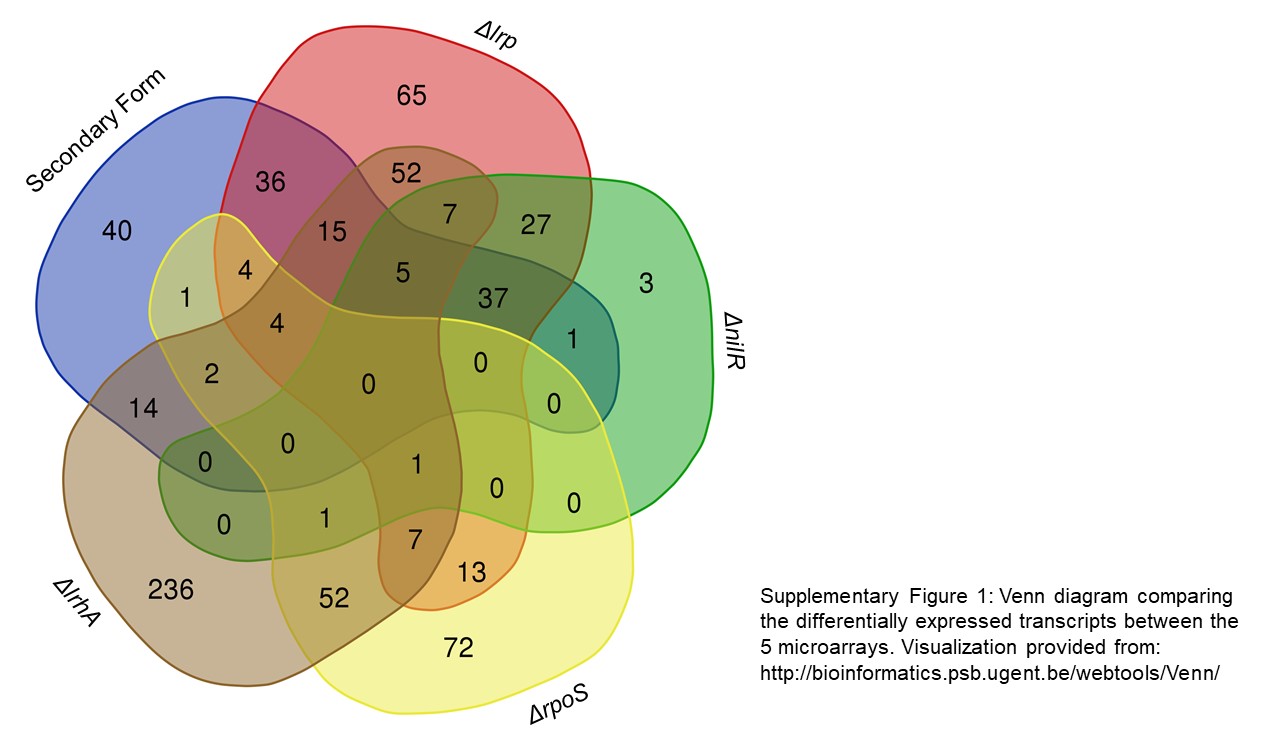

Venn diagram comparing the differentially expressed transcripts between the 5 microarrays analyzed. Visualization provided from: http://bioinformatics.psb.ugent.be/webtools/Venn/

Fig. S2.

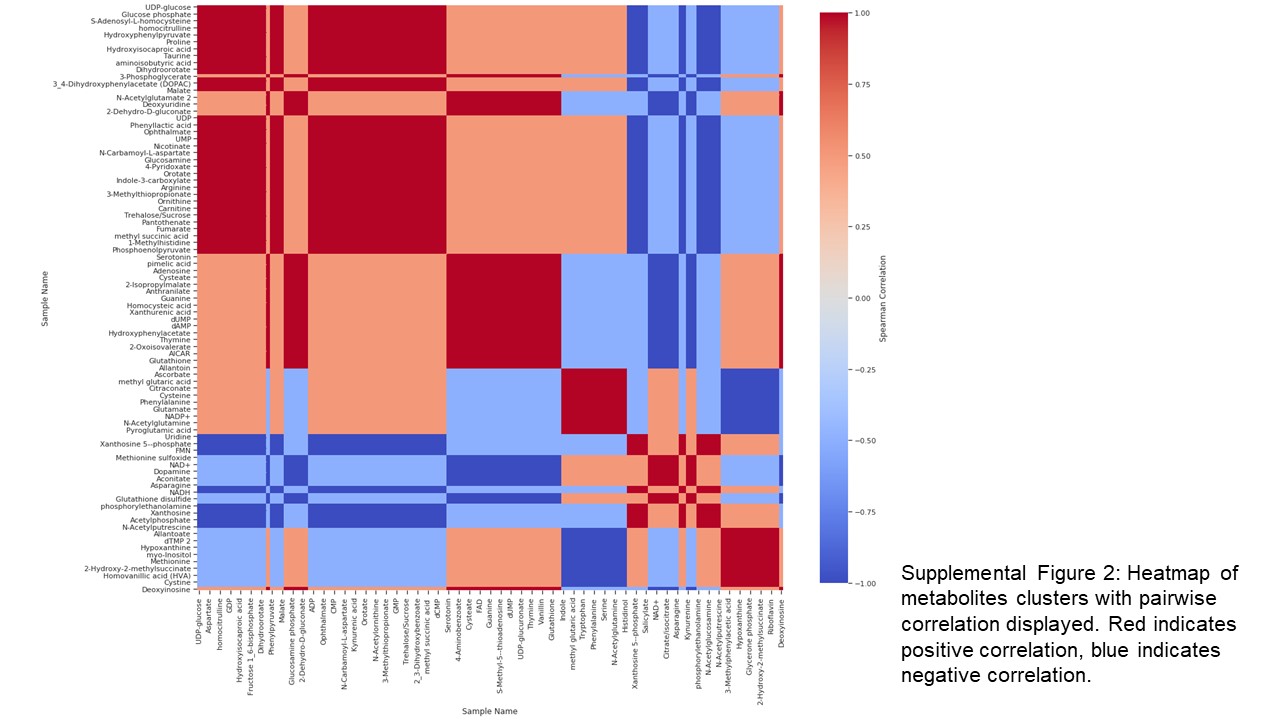

Heatmap of metabolite clusters with pairwise correlation displayed. Red indicates positive correlation, blue indicates negative correlation.

**Fig. S3.**

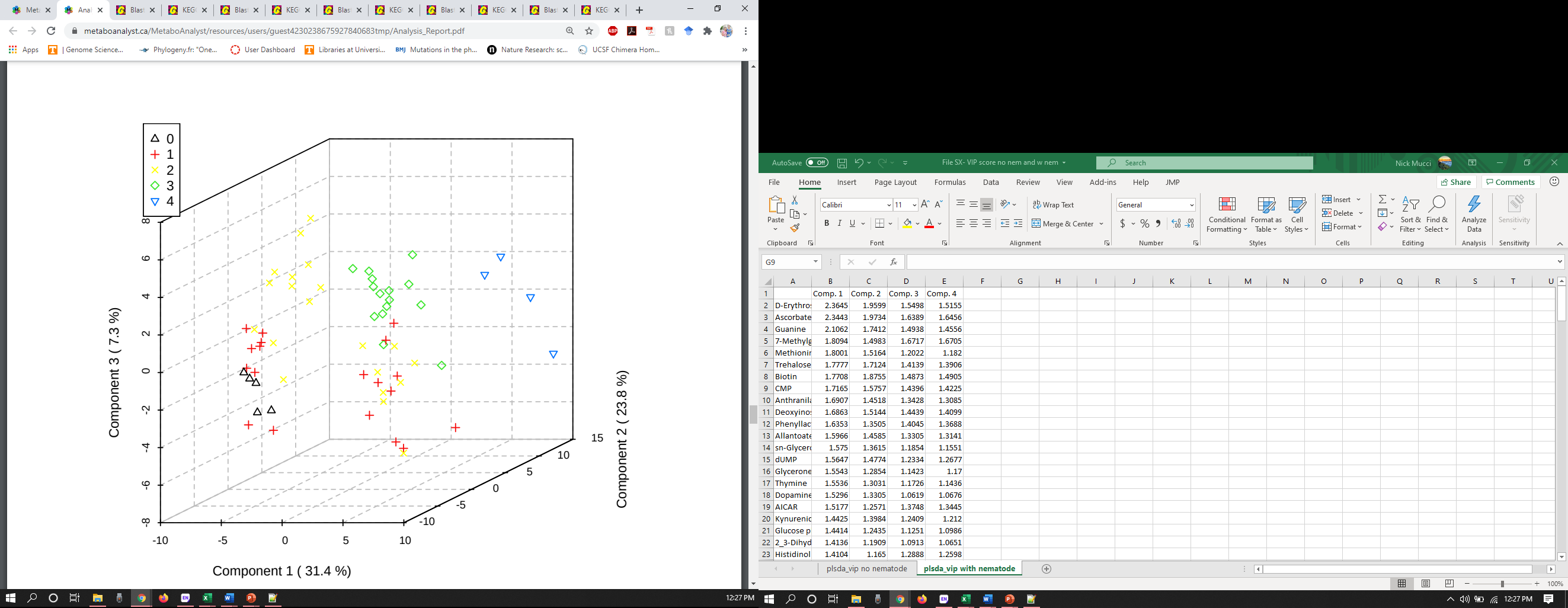

PLS-DA plot for the uninfected insects (black triangles), early phase infected insects (red cross), middle phase infected insects (yellow xs), late phase infected insects (green diamonds), and input nematode IJs (blue upside down triangles). VIP metabolites contributing to the separation of these phases are available at Data S5.

Table S1.

| **Strain** | **Description** | **Reference** |
| --- | --- | --- |
| HGB007 | Amp^r^; *X. nematophila* wild-type ATCC 19061 | ATCC |
| HGB151 | Amp^r^; Kan^r^; Δ*rpoS::kan*; HGB007 | Vivas *et al.*, 2001 |
| HGB800 | Amp^r^; *X. nematophila* wild-type ATCC 19061 | ATCC |
| HGB1059 | Amp^r^; Kan^r^; *lrp-2::kan*; HGB800 | Cowles *et al.*, 2006; Cowles *et al*. 2007 |
| HGB1320 | Amp^r^; Kan^r^; Δ*lrhA2*; HGB800 | Richards and Goodrich-Blair, 2010 |
| HGB1061 | Amp^r^; HGB800 secondary form | Cowles *et al*., 2006 |

List of strains used in the microarray analysis.

Table S2.

| Groups | Un-infected | 1 hour  post-infection | 12 hours alive | 24 hours alive | 24 hours dead | 2 days | 4 days | 6 days | 8 days | 10 days | 12 days | 16  days  (Plate 6) |
| --- | --- | --- | --- | --- | --- | --- | --- | --- | --- | --- | --- | --- |
| Plate 1 | 0.21 | 0.20 | 0.29 | 0.22 | 0.26 | 0.24 | 0.26 | 0.18 | 0.18 | 0.15 | 0.23 | 0.05 |
| Plate 2 | 0.19 | 0.18 | 0.29 | 0.22 | 0.21 | 0.12 | 0.21 | 0.14 | 0.14 | 0.14 | 0.11 | 0.10 |
| Plate 3 | 0.21 | 0.19 | 0.22 | 0.15  (Plate 6) | 0.21 | 0.23 | 0.21 | 0.13 | 0.17 | 0.17 | 0.19 | 0.06 |
| Plate 4 | 0.20 | 0.26 | 0.22 | 0.23 | 0.28 | 0.19 | 0.21 | 0.16 | 0.24 | 0.23 | 0.25 | 0.15 |
| Plate 5 | 0.13 | 0.18 | 0.15 | 0.13  (Plate 6) | 0.13 | 0.16 | 0.19 | 0.15 | 0.13 | 0.14 | 0.20 | 0.20 |

*G. mellonella* weight (g) upon sampling.

Data S1. (separate file)

Trophic study results. Includes raw data, isocline calculations, and trophic level calculations. Also includes notes pertaining to measurements taken.

**Data S2. (separate file)**

Microarray results. Includes every comparison between wild-type (HGB800 or HGB007) and the mutants (described in Table S1). 2<|fold change signal strength| between the two strains are shown, as are the signals for each of these genes.

**Data S3. (separate file)**

Time course metabolomics known data. Includes the raw and normalized data, statistics on the normalized data (including t-test comparisons between uninfected vs. individual time points, t-test comparisons between uninfected vs. defined time phases, fold change differences, average metabolite abundance for each time point among the replicates taken, the standard deviation for the replicates, the standard error for the replicates, the CV for the replicates, and ANOVA results), and a detailed list of the features included in the hierarchical clustering analysis.

**Data S4. (separate file)**

Time course metabolomics unknown data. Includes a list of the unidentified features, statistics on those features (including average metabolite abundance for each time point among the replicates taken, the standard deviation for the replicates, the standard error for the replicates, and the CV for the replicates), and a summary of those measurements.

**Data S5 (separate file)**

VIP metabolites derived from the PLS-DA plots for both the entire time course, and the entire time course with the addition of the input nematode IJs. Data shown includes VIP scores for every detected metabolite and the top three components.
